# From Hummingbird to Elephant: Amyloid Formation in Natural Transthyretin Variants

**DOI:** 10.64898/2026.07.09.737598

**Authors:** Irina Ritsch, Daniel Scholl, Maria Rafiq, Maria A. Martinez-Yamout, Gabriel C. Lander, H. Jane Dyson, Peter E. Wright

## Abstract

Transthyretin (TTR) is a secreted protein associated with cardiac and other amyloid diseases via misfolding. We have previously shown that agitation of human TTR solutions at neutral pH results in aggregation and fibril formation. Here we report that agitation-induced aggregation of TTR from species with very different heart rates (Anna’s hummingbird, hbTTR, and African elephant, aeTTR) differs from that of human TTR (huTTR). Aggregation of hbTTR is slow and favors formation of smaller, fibrillar aggregates, while aeTTR aggregation is rapid and favors larger, more amorphous particles. Spherical, early-stage oligomeric intermediates were found for all variants by mass photometry and electron microscopy. The slow aggregation of hbTTR matches its resistance to denaturation by 8 M urea. The widely different aggregation behavior exhibited by these naturally occurring TTR variants in response to mechanical agitation under close to physiological conditions provides insight into how small sequence differences can contribute to the evolutionary fitness of different animals.

## Introduction

The tetrameric protein transthyretin (TTR) transports thyroid hormone and retinol-binding protein in blood plasma and cerebrospinal fluid (*1*). Aggregation of misfolded TTR results in cardiac and neurodegenerative amyloidosis (ATTR) (*2-4*), which can be further classified into a familial form that is linked to point mutations in the TTR gene, and a typically late-onset spontaneous form that involves wild-type TTR (*5, 6*).

Amyloid formation is initiated by dissociation of the native tetramer (Figure 1A) and subsequent misfolding of the resulting monomers (*7-11*). Recently, we showed that fluid agitation by rapid stirring can substantially accelerate aggregate formation *in vitro* at neutral pH and physiological TTR concentrations (*12*). The agitation assay was designed to mimic the turbulent fluid flow that may occur in the bloodstream (*13, 14*). Accelerated aggregation of wild type TTR and pathogenic variants was also observed in the presence of shear forces generated by agitation or by a pump-based flow system under proteolytic conditions (*15, 16*).

**Figure 1.**
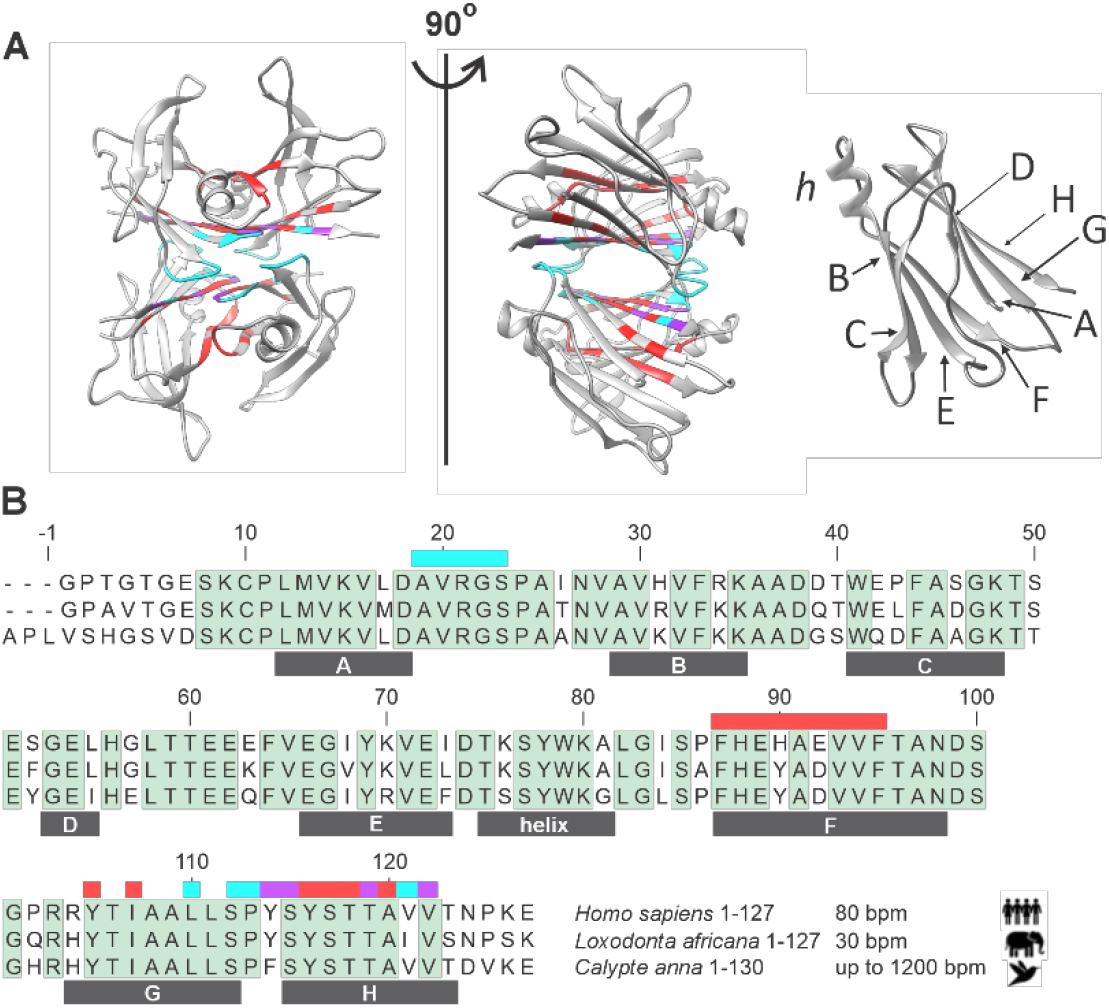
Structure of transthyretin. (A) Left: two orientations of a cartoon model of native human TTR tetramer (PDB 5CN3 (*43*)) colored by subunit interfaces (*44*) (red: strong dimer A-B/A’-B’; blue: weak dimer A-A’/B-B’ and A-B’/A’-B; purple: sites that are involved in both types of interface. Right: cartoon model of a TTR protomer with annotation of the β-strands (A-H), and the EF helix h (PDB: 1TTA (*45*)). (B) Sequence alignment of human WT TTR (huTTR) with those of African elephant (*Loxodonta africana*, aeTTR) and Anna’s hummingbird (*Calypte anna*, hbTTR); the secondary structure elements in huTTR (β-strands A-H and the EF helix) and tetramerization interface residues are shown with colored blocks according to the scheme in part A.

Shear forces from fluid flow are ubiquitous in the vascular system (*17*) and blood flow frequently transitions from laminar flow in the young to more turbulent flow throughout aging (*18*). We hypothesize that if shear forces are an important factor in TTR amyloidosis, then variations in these forces not only with aging, but also across species with highly variable cardiovascular physiology and circulation, might be expected. Species with physiologies that are more prone to high shear forces in the bloodstream may consequently have experienced selective evolutionary pressure for shear-resilient TTR sequences. For example, an increased heart rate is directly correlated with faster blood flow, which is in turn expected to induce more shear stress (*19, 20*). The typical resting heart rate in adult humans is 60-100 beats per minute (bpm), which is towards the lower end of the wide range of resting heart rates across different animal species (Figure S1A) (*21*). Lower resting heart rates (∼30 bpm) are found in African elephants, the largest land-based animals (*22*). Birds in general have higher resting heart rates than mammals, and hummingbirds in particular are among the species with the most extreme variation of heart rate, ranging from < 50 bpm during torpor to around ∼1200 bpm during hover flight (*23*). Given that hummingbirds have a surprisingly long life-span (reported as twelve years in captivity) for their body size (*24, 25*), we hypothesized that hummingbird blood serum proteins, exemplified by TTR, may have evolved to be stabilized against shear stress in blood flow. Conversely, selective pressure on stabilization of African elephant proteins against strong laminar shear stress is expected to have been low, possibly producing a TTR sequence that can poorly resist aggregation under agitation. In this study, we investigated differences in the propensity of TTR derived from these physiological extremes to aggregate under shear forces from rapid stirring (agitation) by applying a previously described experimental approach (*12*).

## Results

### Hummingbird and elephant TTR maintain a native tetrameric state

To identify potential adaptations to the differing blood flow conditions in these species, we compared the amino acid sequences of the TTR from Anna’s hummingbird (*Calypte anna*) (hbTTR), African elephant (*Loxodonta africana*) (aeTTR) and human (*Homo sapiens*) (huTTR) (Table S1). Regions with greater or lesser sequence conservation were identified by aligning these two protein sequences with that of mature human wild-type TTR using blastp (*26*) (Figure 1B) . The alignment of a larger set of sequences corresponding to the heart rate data of Figure S1A is shown in Figure S1B. Of these, hbTTR has the highest sequence identity (95%) with its closest related species, chicken (*Gallus gallus*). The last common ancestor between hummingbird and chicken is estimated to have lived ∼91 Ma ago (https://timetree.org/), which is a similar separation to that between human and African elephant (∼99 Ma). A phylogenetic tree analysis of the selected TTR sequences (Figure S1C), roughly follows the expected species separation. As expected, given the closer relation of the species, the sequence comparison showed that huTTR has higher sequence identity with aeTTR (82%) than with hbTTR (76%). The length of TTR is conserved except for three additional amino acids at the (flexible) N-terminus of hbTTR. Sequence variations occur predominantly in exposed loop regions (Figure 1A, B), the amino- and carboxy termini, and β-strands C, D and E. The latter strands are known hotspot regions for ATTR-associated mutations (*27*). The most conserved regions are the terminal strands A and H, as well as strand G, which is sandwiched between them. Several of the conserved sites are part of the native tetramer interfaces (Figure 1A), but we could confirm that all TTR variants form native tetramers at neutral pH. In analytical size-exclusion chromatography (SEC), hbTTR and aeTTR have very similar elution volumes to huTTR, showing that all three variants are tetrameric (Figure S2). The tetrameric state was dominant in the SEC profiles in the range from ∼0.25-16 μM, and only traces of monomeric or oligomeric species were observed at the extremes of this range. In reverse phase HPLC, aeTTR elutes at higher concentration of organic solvent in the mobile phase than hbTTR and huTTR, which elute similarly (Figure S3C). Based on the theoretical isoelectric point (Table S2), the overall charge of aeTTR in agitation buffer (pH 7) is the closest to neutral and the aeTTR tetramer exhibits the lowest mobility in a native gel assay (reference lane ‘0 h’ in Figure 2A). AeTTR is thus identified as the most hydrophobic variant and closest to neutral charge at pH 7.

**Figure 2.**
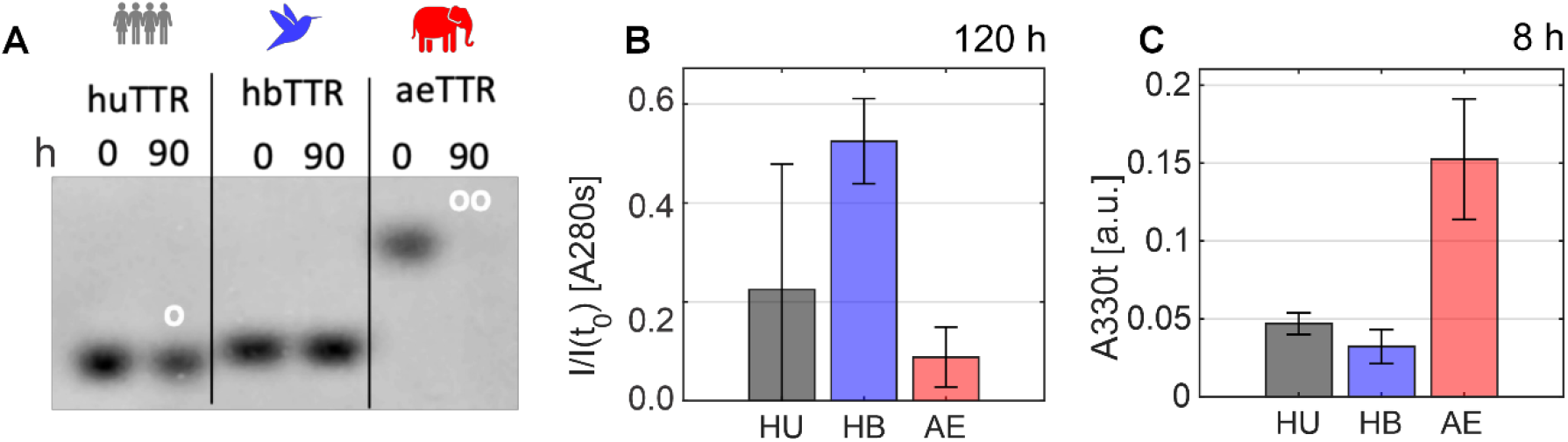
Agitation-induced aggregation assay. A. Native gel electrophoresis of TTR variants (c_0_ = 10 μM) before agitation (0 h) and after 90 hours of agitation, showing depletion of soluble TTR through aggregation. Depleted (o) and strongly depleted (oo) lanes are marked. B. Comparison of supernatant absorbance at 280nm (*A*_280(s)_), corrected for scattering contributions and normalized to the value at t_0_ = 0 h, after 120 h agitation, for human (gray), hummingbird (blue) and elephant (red). (C) Comparison of scattering at 300nm of total sample (*A*_300t_) after 8h. The initial TTR concentrations (*c*_0_) were huTTR, 12.6 μM; hbTTR,13.9 μM; aeTTR,13.3 μM.

### Behavior of hbTTR, huTTR, and aeTTR under agitation

Solutions of huTTR, hbTTR and aeTTR proteins (initial concentration *c*_0_ = 10 μM) were subjected to agitation for 90 h in conical microcentrifuge tubes. The aeTTR samples appeared turbid at the endpoint of the screen, while huTTR and hbTTR samples remained clear. Loss of soluble protein and the amount of tetramer remaining in solution was determined by subjecting the supernatant fraction to native gel electrophoresis. The three proteins behaved differently (Figure 2A). Soluble aeTTR tetramer was almost fully depleted from the supernatant fraction after 90 h agitation, while soluble huTTR tetramer was only moderately depleted and no noticeable change was observed for the hbTTR tetramer. Quantification could be achieved by comparison of the absorbance of the supernatant at 280 nm (*A*_280(s)_), which was corrected (*28*) by subtraction of a weak contribution from traces of residual scattering material remaining after centrifugation (see SI). In Figure 2B we show corrected *A*_280(s)_ for the variants after 120 h agitation (*c*_0_ ∼ 12 μM). In agreement with the native gel result, we found that hbTTR is less susceptible to agitation-induced aggregation than is the case for huTTR and aeTTR. Formation of insoluble aggregates is indicated by increase in the absorbance (scattering) of the total sample before centrifugation at 330 nm, (*A*_330(t)_). Following agitation, *A*_330(t)_ showed the expected increase due to formation of insoluble aggregates, and after as little as 8 h agitation the *A*_330(t)_ for aeTTR was already substantially higher than for the other variants (Figure 2C). The results indicate that these naturally occurring TTR variants differ substantially in their stability towards agitation-induced aggregation at neutral pH.

### Kinetics of hbTTR, huTTR, and aeTTR aggregation

The kinetics of agitation-induced aggregation of each of the TTR orthologs was monitored by measuring light scattering (*A*_330(t)_, *A*_600(t)_), the residual soluble TTR concentration after removal of the insoluble fraction by centrifugation (*A*_280(s)_), and thioflavin T (ThT) fluorescence intensity at various time points for independently agitated samples with initial concentration *c*_0_ in the range of ∼5-20 μM (Figure 3, complete data sets in Figures S4-S7). Kinetic parameters were extracted using a sigmoidal model fit (*12*), modified to include a constant offset parameter for better comparison between variants (Equations S1 and S2), and are summarized in Table S3. Representative data for *A*_330(t)_ and *A*_280(s)_ are shown in Figures 3A and B for samples with *c*_0_ ∼13 μM. The aggregation time course profiles for the three TTR variants differ substantially. Consistent with previous results (*12*), *A*_330(t)_ for huTTR (*c*_0_ = 12.6 μM) shows an initial lag phase followed by an exponential increase that converges to a plateau (Figure 3A, S4) with midpoint time *T*_*m*_ = 18 ± 5 h^-1^. Aggregation of hbTTR (*c*_0_ = 13.9 μM) is much slower, with an extended lag phase (*T*_*m*_ = 77 ± 17 h^-1^), and barely reaches a plateau after 240 h agitation. In contrast, aggregation of aeTTR (*c*_0_ = 13.3 μM) is very rapid without a pronounced lag phase and midpoint time *T*_*m*_ = 5 ± 1 h^-1^. *A*_280(s)_ shows inverted profiles, decreasing with increasing agitation time as soluble protein is lost to aggregates (Figure 3B, S5). Successful removal of large aggregates was confirmed by light scattering measurements with the supernatant samples (*A*_330(s)_, *A*_600(s)_).

**Figure 3.**
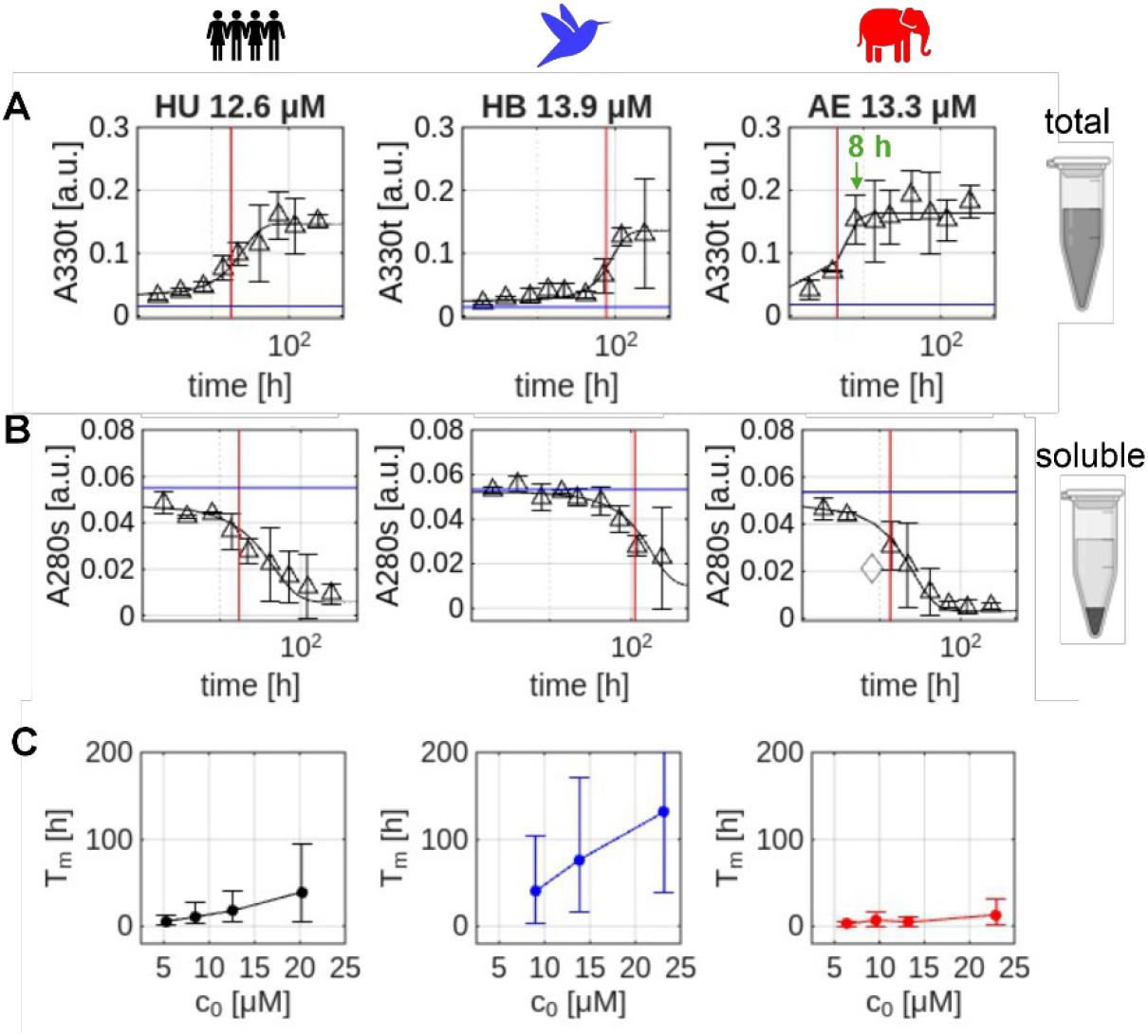
Kinetic modelling of TTR variant aggregation. A, B. Sigmoidal fits of aggregation time courses (solid lines) for TTR variants (c_0_ ∼13 μM; left to right: huTTR, hbTTR, aeTTR). The initial values at t = 0 h are shown as blue horizonal lines; fitted transition midpoint times *T*_*m*_ are marked by red lines. Aggregation profiles for different *c*_0_ values are shown in Figures S4-S7. A. scattering intensity *A*_330(t)_. B. Loss of supernatant protein *A*_280(s)._ The open diamond shows an outlier data point not included in the fit. C. Transition midpoint time *T*_*m*_ for *A*_330(t)_ (Equation S1) versus *c*_0_.

Consistent with a previous report (*12*), we found that higher initial concentration *c*_0_ extended the lag phase, reflected in longer *T*_m_ for huTTR and hbTTR (Figure 3C, Table S3). For aeTTR, *T*_m_ for the *A*_330(t)_ and *A*_600(t)_ scattering profiles was largely independent of *c*_0_ from 6.4-13.3 μM but increased slightly for *c*_0_ = 23 μM. The lag phase for hbTTR was longer and that for aeTTR was shorter than that of huTTR across all concentrations *c*_0_ (Figure S4-S7). The exponential rate *k* followed the opposite trend and was consistently higher for aeTTR than huTTR and lower for hbTTR (Table S3). For huTTR, we previously reported that *T*_m_ for the scattering fits at *A*_*6*00(t)_ is proportional to the expected fraction of monomer *f*_m_ = [M]/*c*_0_, (with *c*_0_ = 4[T] + [M]) which can be numerically determined from the tetramer *K*_D_ and *c*_0._ (*12, 29*). This trend was reproduced by the scattering data in the current work (Figure S8), confirming that the length of the lag phase is related to the availability of free monomer at the start of the experiment. We also observe a positive correlation in a log-log plot of the exponential phase rate *k* as a function of *f*_m_ (Figure S8). The fraction of monomer at the start of the experiment thus not only controls nucleation, which is expected to dominate *T*_m_, but also the rate of aggregate growth in the exponential phase.

### Aggregates from different TTR variants show differences in ThT fluorescence

Kinetic analysis of light scattering data time courses showed that aggregation of hbTTR was slower and aggregation of aeTTR faster than huTTR (Figure 3A, Table S3). Comparison of the scattering values at the two wavelengths (330 nm and 600 nm) with ThT fluorescence gain (Figure S9, Table 1) can give insights into the relative sizes of particles in each sample. For all three TTR variants, a linear correlation is observed between *A*_330(t)_ and *A*_600(t)_ (Figure S9A), but light scattering at 600 nm is disproportionately large for aeTTR and suppressed for hbTTR, suggesting aeTTR forms larger particles. Indeed, aeTTR samples rapidly become cloudy upon agitation. Similarly, the ThT fluorescence intensity gain for each variant is linearly correlated with the extent of aggregation assessed by *A*_330(t)_ and *A*_600(t)_ (Figure S9). To monitor the extent of soluble protein loss, we converted the residual *A*_280(s)_ for all time points and concentrations *c*_0_ back to a supernatant molar protomer concentration *c*_s_, and subtracted *c*_*s*_ from *c*_0_ (*c*_*0*_ – *c*_s_, in μM).

**Table 1.** Cross-parameter correlation analysis. The linear regression slope and R^2^ of ThT fluorescence intensity gain versus loss of supernatant TTR (c_0_-c_s_, μM) was derived from replicate averaged data (Figure 4A).The fitted parameters for ThT gain versus *A*_600(t)_, ThT gain versus *A*_330(t)_, and *A*_600(t)_ versus *A*_330(t)_ were obtained from the data for all replicates (Figure S9).

| Variant | ThT gain<br>vs ($c_0 - c_s$) | | ThT gain<br>vs $A_{600(t)}$ | | ThT gain<br>vs $A_{330(t)}$ | | $A_{600(t)}$<br>vs $A_{330(t)}$ | |
| --- | --- | --- | --- | --- | --- | --- | --- | --- |
| | slope | $R^2$ | slope | $R^2$ | slope | $R^2$ | slope | $R^2$ |
| HU | $3.7 \pm 0.2$ | 0.90 | $600 \pm 20$ | 0.90 | $230 \pm 10$ | 0.80 | $0.39 \pm 0.01$ | 0.95 |
| HB | $7.5 \pm 0.9$ | 0.76 | $1330 \pm 50$ | 0.92 | $460 \pm 20$ | 0.89 | $0.34 \pm 0.01$ | 0.94 |
| AE | $2.9 \pm 0.4$ | 0.70 | $400 \pm 30$ | 0.73 | $170 \pm 20$ | 0.62 | $0.45 \pm 0.01$ | 0.97 |

For all three variants, a linear correlation was observed between the ThT fluorescence intensity gain and (*c*_*0*_ – *c*_s_) (Figure 4A). The slope of the correlation between ThT fluorescence gain and loss of soluble TTR (*c*_*0*_ – *c*_s_) was more than two times higher for hbTTR than for aeTTR, with huTTR forming the intermediate case (Table 1). Analogous trends can be found for linear regressions between *A*_330(t)_ or *A*_600(t)_ and ThT gain (Table 1), which directly correlate the amount of scattering material to the amount of fibrillar material (Figure S9). For a given *c*_*0*_, the aggregates formed by hbTTR had higher fibrillar content than those of aeTTR, despite slower aggregation kinetics than aeTTR, which yielded greater amounts of aggregated material at faster rates. Further insights into the formation of fibrillar material could be gained from the fits of the ThT fluorescence intensity gain time courses (Figure S7, Table S3). For huTTR, the fitted parameters are the same within experimental uncertainties for the ThT, *A*_330(t)_, and *A*_600(t)_ time course profiles, but because aggregation of hbTTR was very slow the ThT fluorescence intensity gain did not reach a plateau within our dataset, so the fitted parameters are imprecise. Overall, the plateaus in the ThT gain graphs for aeTTR were lower, but at higher *c*_0_ (13.3 and 23 μM), the ThT fluorescence intensity gain for aeTTR appeared to be biphasic (Figure S7), with a long *T*_m_ process that affects ThT fluorescence superimposed on the short lag phase aggregation observed at lower concentration. Sigmoidal fits restricted to the early time points, omitting the 240 h data point, yielded *T*_m_ values similar to those determined from the scattering data (Table S3, Figure S7).

**Figure 4.**
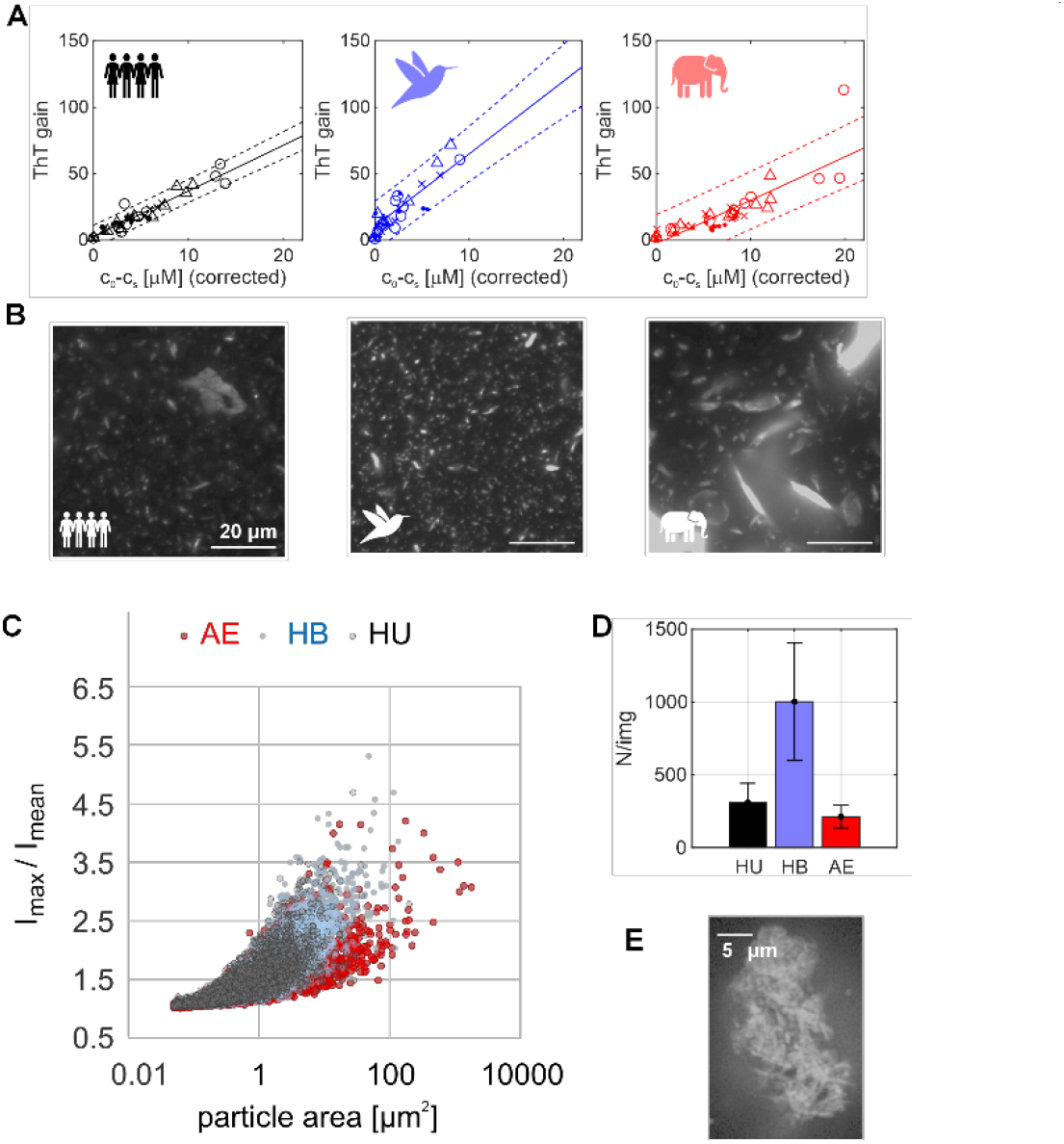
Characterization of transthyretin particles. A. Combined linear regression (solid line, dotted lines show 95% confidence interval) of loss of supernatant TTR (*c*_0_-*c*_s_) versus ThT fluorescence intensity gain across the aggregation time course for *c*_0_ ∼5 μM (solid dots), ∼9 μM (x), ∼12 μM (triangles), and ∼20 μM (open circles). Datapoints with less than 5% or more than 95% loss of *c*_0_ were omitted from the fit. Linear regression parameters are listed in Table 1. B. ThT fluorescence microscopy (100x magnification, images displayed slightly oversaturated) of samples after 74h agitation (*c*_0_ ∼10 μM, 200 μM ThT added post-agitation); C, D. Particle statistics from image series in panel B, using a thresholding analysis pipeline (see Table 2). C. Correlation of particle size with intensity variation within each particle (maximum intensity in particle boundary I_max_ and mean intensity in particle I_mean_). D. Number of particles per image. E. 100x image of a huTTR aggregate with resolution increased by 1D pseudo-confocal filtering.

### Imaging reveals different aggregate morphologies

To investigate whether hbTTR and aeTTR aggregates differ in their morphology (fibrillar versus amorphous), we imaged ThT-stained aggregates by fluorescence microscopy. Representative images of aggregates from agitated samples (*c*_0_ = 10 μM, 74h agitation, n=2 replicates per variant) are shown in Figure 4B and Figure S10. For this choice of *c*_0_ and time point, huTTR and hbTTR had intermediate and aeTTR substantially higher loss of supernatant protein (normalized c_s_-c_0_, given as percentage of c_0_ in Table 2). For imaging, a total aliquot of agitated sample was mixed with ThT (final concentration 100 μM) and aggregates were left to sediment onto a glass coverslip under quiescent conditions for ∼1 h following agitation. By imaging sedimented particles on the glass we can be confident that illumination of the aggregates was as consistent as possible across the sample well, although this may introduce a small bias in particle statistics due to differences in size- and shape-dependent sedimentation rates, as well as different adherence to the weakly negatively charged silicate glass surface. ThT positive aggregates were observed in all cases; hbTTR showed many small particles (∼1 μm^2^ cross-section) whereas aeTTR samples showed fewer and substantially larger particles (> 20 μm^2^). Image stacks perpendicular to the glass surface (z-stacks) revealed that some of the hbTTR aggregates that appear to have very small cross-sections (< 1 μm^2^) in individual focal planes are in fact thin, elongated aggregates that remain oriented perpendicular to the glass surface (Figure S11) despite the long sedimentation time, but videos captured directly above the glass surface showed comparatively faster Brownian motion of hbTTR aggregates consistent with overall smaller objects (Supporting Videos). Quantitative comparisons were made by calculating average aggregate numbers and size statistics in a series of images from multiple replicates using thresholding workflows in ImageJ (https://imagej.net/software/imagej/). Examples of the image analysis are shown in Figure S10 and S12. The particle statistics confirm that aggregates formed by aeTTR were substantially larger (particle area *A*_mean_ in Table 2), and with more size variation, than those of huTTR or hbTTR (Figure 4C, S12). The mean ThT fluorescence intensity within each identified particle (*I*_mean_(particle), Table 2) indicates a similar ThT binding capacity for huTTR and hbTTR aggregates (Figure S12). *I*_mean_ was smaller for the aeTTR aggregates, suggesting a lower β-strand content and a higher proportion of amorphous material (Figure S12C). We can calculate the fluorescence intensity of an average particle for each variant (*I*_particle_ = *I*_mean (particle)_ x *A*_mean (particle)_), and multiply by the average number of particles per image *N*_av_ (same images sizes were used consistently) to obtain an estimate of the bulk fluorescence measurement (*I*_sample_= *I*_particle_ x *N*_av_). The estimate from *I*_sample_ recapitulated the trends of the bulk measurement well (ThT gain in Table 2), adding confidence to the particle analysis. In combination, the unexpectedly high ratio of bulk ThT fluorescence intensity gain for hbTTR relative to light scattering at 600nm (Figure S9C, Table 1) can be attributed to a high count of very small particles (Figure 4D) that likely have smaller light scattering cross-sections than the larger or differently shaped aggregates formed by other variants. For huTTR, we acquired high-resolution fluorescence images using a pseudo-confocal, Fourier filtering-based imaging technique, which revealed that at least a fraction of the larger aggregates in huTTR were internally composed of bundles of elongated, fibrillar components (Figure 4E), highlighting that aggregate morphology and maturation must be considered as a parameter when comparing the aggregation kinetics of TTR variants in bulk assays.

**Table 2.** Particle statistics from ThT fluorescent imaging (Figure 4B and Figure S10-S12) using two independently agitated replicates per variant (c_0_ =10 μM, 74 h agitation). Images (100x magnification, 2048×2048 pixel,) were acquired at ten randomly selected locations per sample well and processed as a combined image stack (total N_image_ =19 in aeTTR and hbTTR stacks, 18 in huTTR stack, after removal of images at the well edge; error estimates represent standard deviations between images in the stacks). The columns report the average number of particles per image N_av_, the density of particles ρ per image area (133 x 133 μm^2^/image), the mean particle cross-section area *A*_mean_, and the average ThT fluorescence intensity per particle *I*_mean_(particle). The contribution of an average particle to the bulk fluorescence signal was estimated as *I*_particle_ = *I*_mean_ (particle) x *A*_mean_(particle) and used to estimate the combined ThT intensity per image, *I*_sample_ = N_av_ x *I*_particle_. The experimental bulk ThT fluorescence intensity gain and scattering at 330 nm were measured with a standard plate reader assay on the same samples and averaged over the two replicates. Loss of supernatant protein relative to initial (c_s-_c_0_)/c_0_ was tabulated based on replicate-averaged nanodrop measurements.

| Variant | $N_{\text{av}}$ | $\rho$ [$1/\mu\text{m}^2$] | $A_{\text{mean}}$ (particle) [$\mu\text{m}^2$] | $I_{\text{mean}}$ (particle) [cts] | $I_{\text{particle}}$ [cts] $\times 10^3$ | $I_{\text{sample}}$ [cts/image] $\times 10^6$ | ThT gain | A330t [a.u.] | norm. $c_s - c_0$ [% initial] |
| --- | --- | --- | --- | --- | --- | --- | --- | --- | --- |
| HU | $300 \pm 130$ | $0.02 \pm 0.01$ | $1.0 \pm 0.3$ | $4100 \pm 100$ | $4 \pm 1$ | $1.0 \pm 0.7$ | $24 \pm 5$ | $0.11 \pm 0.02$ | $40 \pm 20$ |
| HB | $1000 \pm 400$ | $0.06 \pm 0.02$ | $1.4 \pm 0.3$ | $4200 \pm 30$ | $6 \pm 1$ | $6 \pm 3$ | $60 \pm 10$ | $0.11 \pm 0.03$ | $45 \pm 1$ |
| AE | $210 \pm 80$ | $0.013 \pm 0.004$ | $5 \pm 4$ | $4000 \pm 100$ | $20 \pm 16$ | $4 \pm 4$ | $40 \pm 15$ | $0.14 \pm 0.05$ | $83 \pm 3$ |

### Tetrameric TTR in the supernatant

Aggregation of TTR into canonical amyloid fibrils requires dissociation of the tetramer and misfolding/unfolding of the resultant monomer (*7, 12, 30*). The extent to which a variant forms fibrils rather than amorphous aggregates is linked to both tetramer stability and monomer unfolding. To assess if there are differences in the tetrameric fraction in the residual supernatant TTR following agitation, a resveratrol (RV) binding assay was performed (for c_0_ > 12 μM only, due to sensitivity limits). RV undergoes a large increase in fluorescence intensity upon binding to the TTR tetramer ligand-binding pocket, which is absent from monomeric TTR (*31*). Addition of RV to quiescent reference samples of all TTR variants led to an increase in RV fluorescence intensity relative to RV in buffer (RV gain > 1, Figure S13). The absolute value of the initial RV fluorescence intensity gain depends on the applied excess of RV over TTR tetramer, which was ∼ 11-fold over the initial tetramer concentration for c_0_ ∼ 12 μM, and ∼ 7-fold for c_0_ ∼ 20 μM. The RV fluorescence intensity gain for unperturbed huTTR was smaller than for hbTTR and aeTTR (Figure S13), which is likely due to intrinsic differences in the fluorescent behavior of the dye when bound to the different TTR binding pockets or differences in site occupancy. All variants showed the expected decrease in RV fluorescence intensity with increasing agitation time (Figure S13).

### Negative stain TEM and mass photometry with oligomers

Characterization of early-stage oligomeric intermediates formed during TTR aggregation, which are always present at very low populations, has been a long-standing challenge. We used a combination of mass photometry and negative stain transmission electron microscopy (TEM), which are sensitive to particles between the size of the native tetramer (∼55 kDa) and the lower size limit of fluorescence microscopy (∼1 μm), to characterize oligomers formed after agitation of TTR solutions. Based on our previous characterization of the aggregation kinetics, we aimed to maximize any potentially oligomeric fraction by using a low starting concentration (*c*_0_ = 2.5 μM) that favors rapid aggregation (short *T*_m_, see Table S3) and applied a short agitation time (24 h) to capture particles before they can mature into larger aggregates. The success of the strategy was confirmed by mass photometry acquired at 50 nM protomer concentration after rapid dilution in the sample well immediately prior to data acquisition. In quiescent reference samples we observed the expected tetrameric mass peak for all variants (number of protomers *n* = 4) with similar tetramer peak mass centers (huTTR: 54 kDa, hbTTR: 47 kDa, aeTTR: 57 kDa, Figure S14A). The tetramers were observed within the time of mass photometry acquisition (one minute acquisition per run, ∼5s mixing time before acquisition), emphasizing the stability of the tetramer assembly of all three natural variants even at very low concentrations. In the agitated samples we observed a shift of the intensity distribution towards the higher molecular weight region, corresponding to formation of higher order oligomers (Figure 5A). We found shoulders on the higher mass flank of the tetramer peak corresponding to a range of up to n ∼ 8 for huTTR (Figure 5A). Even higher order assemblies were sporadically encountered, but counts were too low for reliable quantification. A weak shoulder was also observed on the tetramer peak for hbTTR, but the sample remained predominantly tetrameric. The tetramer fraction was highly depleted in aeTTR, and a long tail of higher MW oligomers was observed, with masses up to at least *n* = 16. No single molecular weight multiple of the protomer clearly dominated, indicating variable stoichiometry of the oligomeric assemblies.

**Figure 5.**
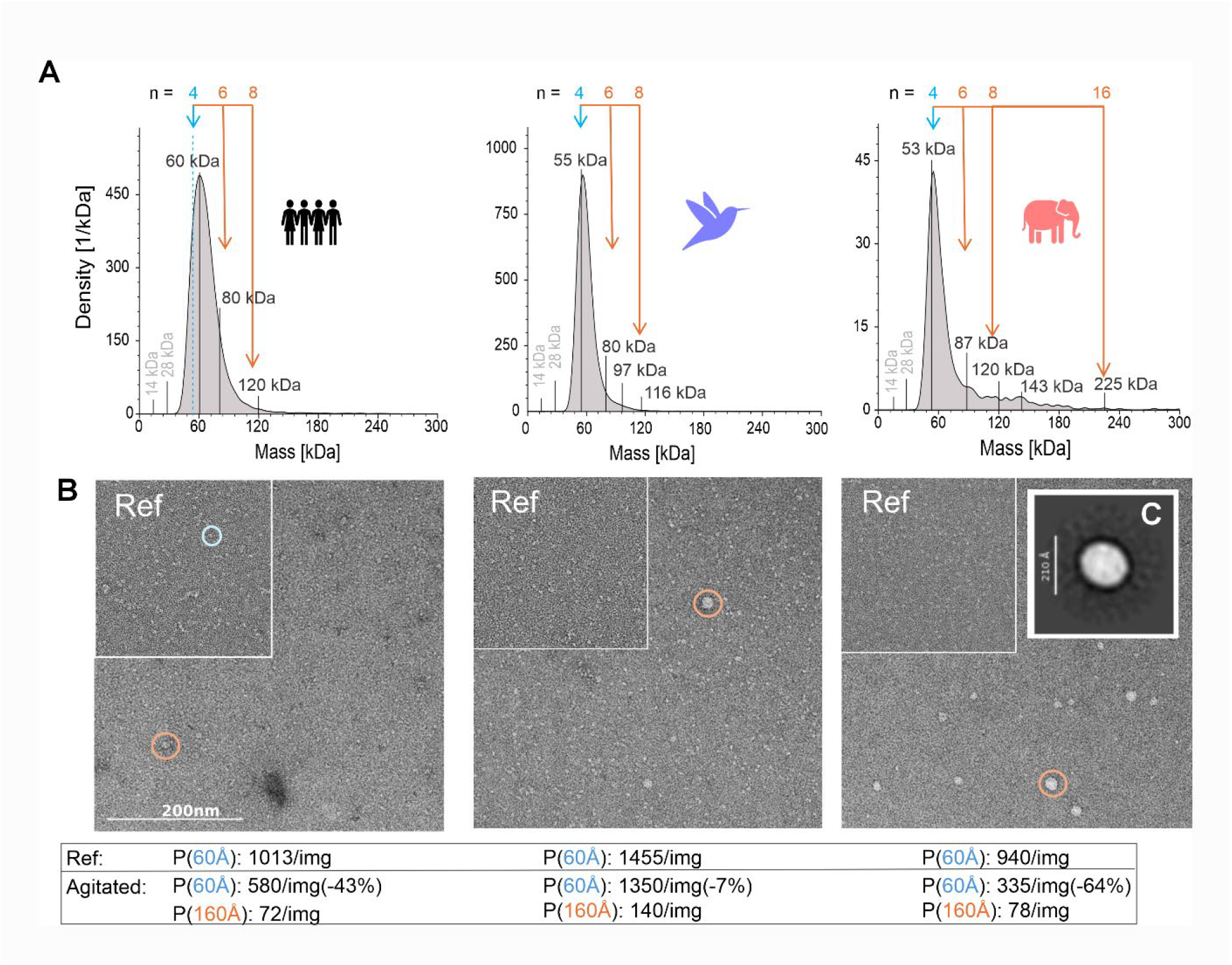
Particle analysis by mass photometry and TEM. A. Mass photometry (kernel density estimation KDE) after 24 h sample agitation (*c*_0_ = 2.5 μM) at room temperature for huTTR, hbTTR and aeTTR. Data were recorded after rapid in-well dilution to 50 nM protomer (1:40 dilution). The expected molecular weights for the tetramer (n = 4) and higher order oligomers are indicated. B. Negative stain TEM of the same samples before (insets labeled Ref) and after 24 h agitation; the blue circle marks one example of a tetrameric particle (< 60Å diameter) and the orange circles mark examples of oligomeric particles. The DoG picker statistics P for the 60 Å particles (range 0.7-1.5) are indicated, the values for the reference were scaled up by a factor 1.25 to account for TTR concentration changes during grid loading (2 μM for Ref, 2.5 μM for agitated). The DoG picker statistics for 160 Å particles (range 0.5-1.75) are indicated for the agitated samples. C. Image of the most populated single particle 2D class (157 of 2059 particles (7.6%) from 74 micrographs) in the agitated sample of aeTTR after large blob picking (200-500 Å) and filtering using cryosparc (scale bar 210 Å).

Negative stain TEM images for the same samples before and after agitation are shown in Figure 5B. In TEM images of reference samples (not agitated) we observed isolated particles of the expected size for TTR tetramers for all three variants (one example is marked with a blue circle in the reference inset in Figure 5B). After 24 h agitation. some tetrameric particles could still be observed, but at substantially lower counts for huTTR (-43% of original counts) and especially aeTTR (-64%). In contrast, tetramer counts in the hbTTR sample remained very high (-7%). TEM and mass photometry are individually ill-suited for quantification of particle statistics due to sampling issues, but in combination they reported consistent results for depletion of tetramer and formation of oligomers across variants. All agitated samples, especially aeTTR, contained particles of sizes larger than tetramer. However, the TEM images of hbTTR contained more of these particles than would be expected from the mass photometry data (Figure 5A). This may result from dissociation of hbTTR oligomers during the ∼40-fold dilution required for mass photometry acquisition. Given the low *c*_0_ of this screen, no dilution was required to prepare the TEM grids, which reduces the risk of oligomer dissolution. Most strikingly, we observed particles with a diameter of ∼15-30 nm in TEM images recorded following 24 h agitation that were spherically symmetric and densely packed, with no clearly discernible internal structure at negative stain resolution (Figure 5B). These spherical oligomers are not an artifact from the acidity of the uranyl formate stain since they were also observed with neutral pH tungsten-based stain (Figure S14B). Uranyl formate gave better contrast and was thus used for the remaining analysis. The oligomeric particles that appeared in TEM images of samples agitated for 24 h were absent from images recorded on samples agitated for a longer 72 h period (Figure S14C) suggesting that they are on-pathway intermediates in the aggregation cascade. The oligomeric particles formed by agitation of aeTTR were abundant and comparatively homogenous in size, which allowed 2D classification. The highest populated class average is shown in Figure 5C and the complete set of classes is shown in Figure S14D. The images did not reveal internal structure, suggesting a highly dynamic assembly. The appearance of these oligomers is reminiscent of the early stages of acid-induced aggregation (*32*).

### Stability of TTR variants

The preceding experiments suggested that the intrinsic tetramer stability of the TTR variants is likely very different, which could explain the differences in aggregation kinetics. Kinetic stability of the tetramer and thermodynamic stability of the TTR protomer can be probed by chemical denaturation in urea using a well-established native Trp fluorescence screen (*31*). The experiment monitors the change in intensity ratio R_355/335_ throughout unfolding based on the excitation spectrum of unfolded versus folded TTR. We confirmed that the intensity ratio methodology can be applied to the hbTTR and aeTTR variants by observing the expected folded ratio (*R*_355/335_ ≈ 0.85) at zero denaturant, and unfolded ratio (*R*_355/335_ ≈ 1.4) after unfolding by boiling in 6 M urea for 10 min (Figure S15A). Urea denaturation profiles were measured following 120 h of incubation at ambient temperature in variable concentrations of urea (∼ 0-8 M) (Figure 6).

**Figure 6.**
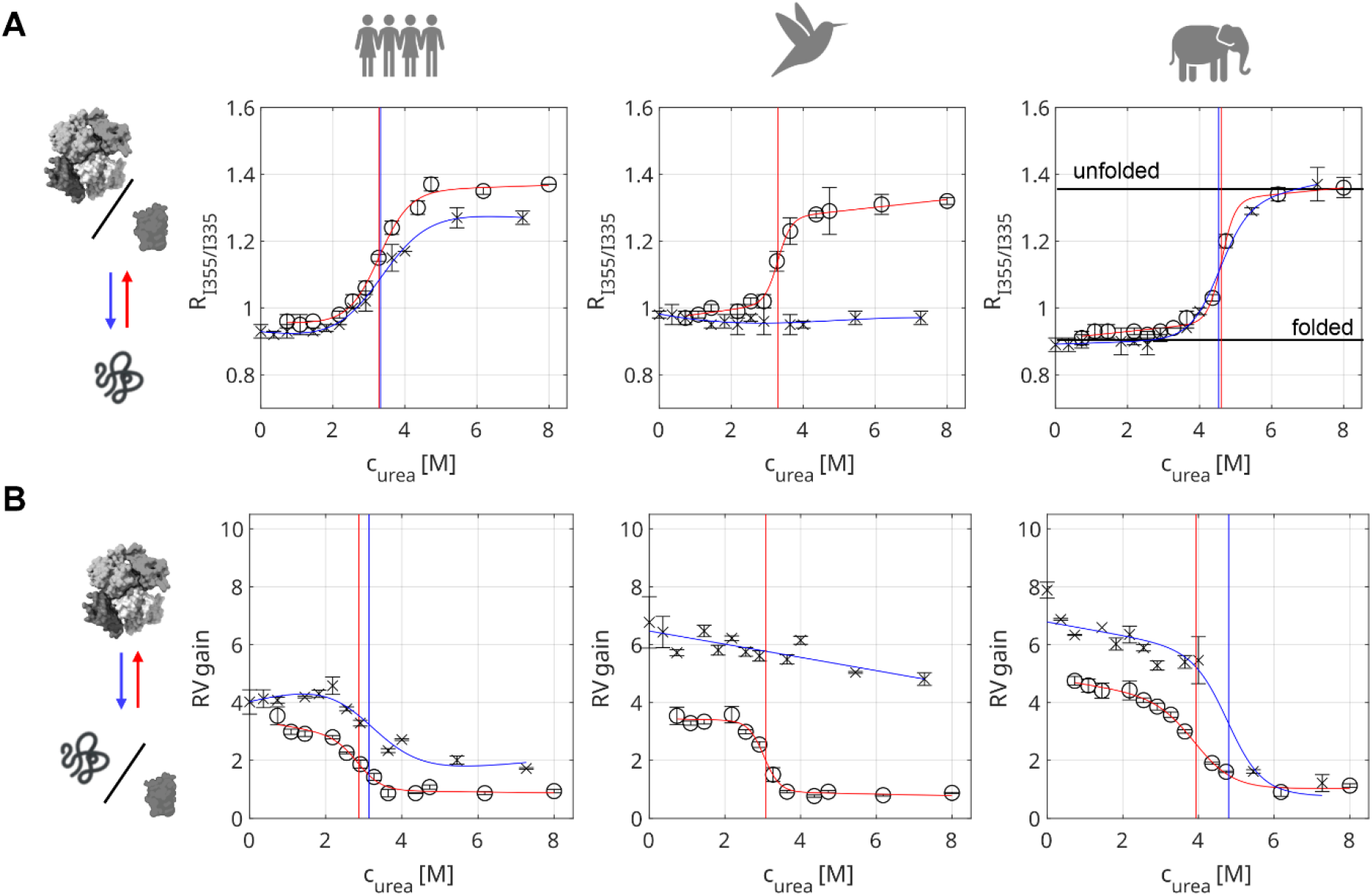
Urea unfolding (x, blue lines) and refolding (o, red lines) assays monitored by A. the Trp fluorescence intensity ratio R_355/335_, and B. the RV fluorescence intensity gain. The fitted transition midpoint urea concentrations (*c*_m_) are indicated by dashed vertical lines. The TTR concentration was 10 μM for all variants and plates were incubated at ambient temperature for 120h.

Both huTTR and aeTTR showed characteristic denaturation profiles with a single defined unfolding transition, whereas after the same time period hbTTR showed no signs of unfolding (Figure 6A), implying that it is kinetically even more stable than the most stable reported human TTR variant (T119M) (*31*). The urea denaturation profiles of huTTR and aeTTR were fitted using the two-state model of Hurshman Babbes et al. (*33*) (Equation S3, blue curves in Figure 6A). The unfolding transition mid-point urea concentration *c*_m_(unfold) is a measure of thermodynamic stability in the folding/unfolding equilibrium, with higher values indicating greater stability. For 10 μM huTTR in agitation buffer we determined *c*_m_(unfold)= 3.3 ± 0.4 M (Table 3), which is in good agreement with literature values (3.4 M urea for 7.2 μM TTR (*33*)). A higher transition mid-point was observed for 10 μM aeTTR (*c*_m_(unfold)= 4.7 ± 0.4 M). In the case of aeTTR and huTTR the transition midpoint *c*_m_(unfold) matched the tetramer dissociation equilibrium mid-point *c*_m,RV_(unfold) observed by RV binding (Table 3, blue curves in Figure 6B), indicating that dissociation is tightly coupled to unfolding. For the forward unfolding reaction, we could also determine kinetic information for huTTR and aeTTR (Figure S15, Table 3).

**Table 3.** Thermodynamic and kinetic stability parameters (fits from Equations S3-S6) from the R_355/335_ urea denaturation for unfolding/refolding and the resveratrol (RV) binding assay for dissociation/reassociation shown in Figure 6. The fit results are referred to as either ‘unfold’ or ‘refold’ depending on which dataset was fitted. The columns show the transition mid-point urea concentration c_m_, the urea dependence factor for unfolding *m*, the transition mid-point for tetramer formation from the RV assay c_m,RV_, the urea dependence factor for dissociation from the RV screen *m*_RV_, and the extrapolated tetramer dissociation rate constant at 0 M urea k_u,0_ and urea dependence of tetramer dissociation rate constant *m*_u_ for huTTR and aeTTR (from fits in Figure S15). Parameters for unfolding/dissociation of hbTTR could not be determined since no change was observed within the limits of the screen.

| variant | $c_m$<br>[M urea] | $m$<br>[kJ/(mol (M urea))] | $c_{m,RV}$<br>[M urea] | $m_{RV}$<br>[kJ/(mol M urea)] | $k_{u,0}$ [1/h] | $m_u$ [1/ M urea] |
| --- | --- | --- | --- | --- | --- | --- |
| unfolding conditions (dissociation) |  |  |  |  |  |  |
| huTTR | $3.3 \pm 0.4$ | $-4 \pm 2$ | $3.1 \pm 0.4$ | $-4 \pm 2$ | $0.042 \pm 0.003$ | $0.08 \pm 0.01$ |
| aeTTR | $4.7 \pm 0.4$ | $-6 \pm 6$ | $4.8 \pm 0.8$ | $-5 \pm 4$ | $0.005 \pm 0.005$ | $0.5 \pm 0.1$ |
| refolding conditions (reassociation) |  |  |  |  |  |  |
| huTTR | $3.3 \pm 0.4$ | $-5 \pm 2$ | $2.9 \pm 0.2$ | $-8 \pm 5$ | | |
| hbTTR | $3.3 \pm 0.1$ | $-9 \pm 4$ | $3.0 \pm 0.2$ | $-12 \pm 7$ | | |
| aeTTR | $4.6 \pm 0.1$ | $-11 \pm 4$ | $4.0 \pm 0.1$ | $-4.8 \pm 0.9$ | | |

Unfolding of TTR at urea concentrations above *c*_m_(unfold) is rate-limited by tetramer dissociation, which allows extrapolation of the unfolding rates at high urea to zero molar denaturant as an estimate for intrinsic tetramer dissociation rate *k*_u,0_, with urea dependence factor *m*_u_ (*31*). The extrapolated *k*_u,0_ of aeTTR was approximately eight times smaller than for huTTR, while the urea dependence *m*_u_ was almost six times larger (Table 3).

Kinetic analysis was not possible for hbTTR due to insufficient signal change over the assay time. To access thermodynamic stability parameters of hbTTR, we applied a refolding approach. All variants, including hbTTR, could be denatured by incubation in 6 M guanidinium hydrochloride (GdnHCl) for 48 h. Samples that were thus unfolded were allowed to refold in a series of urea buffers (red curves in Figure 6A,B). All variants were capable of refolding and the process was approximately complete within 24 h (Figure S15) but was allowed to continue for 120 h for consistency with the unfolding reaction. The *c*_m_(refold) values for the refolding transition for huTTR and aeTTR matched those for the unfolding transition (Table 3). In contrast, pronounced hysteresis was observed in the unfolding and refolding transitions of hbTTR (Figure 6A), which refolded with a c_m_(refold) of 3.3± 0.1 M while being resistant against unfolding at urea concentrations up to 8 M. After refolding, all three variants were able to bind RV, with a reassociation transition that matched the refolding transition (*c*_m,RV_(refold) in Table 3) but the RV fluorescence gain after refolding was consistently lower than for samples that had never undergone an unfolding/refolding cycle, likely due to incomplete formation of natively folded tetramer. In summary, we found that hbTTR dissociation was very slow under urea denaturation conditions, but refolding from the unfolded state proceeded with similar efficiency as for huTTR, indicating that the protomer fold stability is similar. The protomer stability of aeTTR was higher than huTTR both in unfolding and refolding processes.

## Discussion

The present work provides insights into the aggregation propensities of natural TTR variants, from species with widely varying heart rates, when exposed to chaotic shear forces associated with stirring in microcentrifuge tubes at neutral pH. At comparable initial concentrations (*c*_0_ ∼13 μM), agitation-induced aggregation of TTR from African elephant (resting heart rate ∼30 bpm) is very rapid (*T*_m_ = 5 h by light scattering) while aggregation of TTR from hummingbird (heart rate ∼1200 bpm during hover flight) is very slow (*T*_m_ ∼80 h) and barely reaches the plateau phase after 240 h agitation (Figures S4, S5). Aggregation of human TTR (resting heart rate 60-100 bpm) occurs at an intermediate rate (*T*_m_ ∼20 h). These observations suggest that hummingbird TTR, which is likely exposed to high shearing forces at peak blood flow rate during hover, may have evolved to be stable against shear-induced aggregation. Interestingly, TTR from the mouse (heart rate 500-700 bpm) is also kinetically stable and is non-amyloidogenic (*34*). In contrast, evolutionary pressures to stabilize TTR against shear forces are expected to be low for African elephants, with low resting heart rates, consistent with the rapid aggregation of aeTTR upon agitation.

In accord with previous observations (*12*), the lag phase for agitation-induced aggregation of huTTR becomes longer as the total tetramer concentration *c*_0_ is increased (Figures S4-S7, Table S3). Both *T*_m_ and the exponential aggregation rate *k* are dependent on the fraction of monomer *f*_m_, with *T*_m_ decreasing and *k* increasing as the fraction of monomer increases (Figure S8). This behavior is consistent with a mechanism in which the huTTR tetramer must first dissociate to monomer before entry into the aggregation pathway (*12*). The hbTTR behaves similarly to the human protein, with *T*_m_ increasing and *k* decreasing as the total tetramer concentration increases, showing that agitation-induced aggregation occurs by the same dissociative pathway as huTTR. In contrast to huTTR and hbTTR, *T*_m_ for aeTTR is independent of total tetramer concentration from 6.4 to 13.3 μM (Figures S4-S7, Table S3). This suggests that the initial stages of aeTTR aggregation may proceed by a different mechanism that does not require tetramer dissociation to monomer, or if dissociation does occur, reassembly of the tetramer cannot outcompete downstream aggregation in this concentration range. The aeTTR is only weakly charged at pH 7 (Table S2) and has a propensity to form heterogeneous oligomers and large oligomeric particles visible by TEM after 24 h agitation (Figures 5, S14). The ratio of ThT fluorescence intensity gain versus *A*_330(t)_ and *A*_600(t)_ is smaller than for huTTR and hbTTR (Table1 and Figure S9), suggesting that the oligomeric particles of aeTTR, although efficient at scattering light, have a smaller β-sheet content. The ThT fluorescence time course profiles for aeTTR at 13.3 and 23 μM appear to be biphasic, with a substantial increase in fluorescence intensity at the longest time points (Figure S7), suggesting slow progression of the initially formed oligomers towards more β-rich structures.

In addition to dissociation of the TTR tetramer, unfolding and structural remodeling of the monomer are a prerequisite for amyloid fibril formation (*8, 9, 35*). Previous studies have shown that the amyloidogenicity of pathogenic human TTR variants is related to the thermodynamic and kinetic stabilities of the tetramer (*3, 31*). We therefore performed urea denaturation/renaturation experiments to probe the relative stabilities of huTTR, hbTTR and aeTTR. The hbTTR tetramer showed no detectable dissociation or unfolding at urea concentrations up to 8 M (Figure 6) and, consistent with this, was relatively resistant to agitation-induced aggregation, exhibiting a much longer lag phase than huTTR at similar starting concentrations (Figures S4-S7, Table S3). Although the hbTTR tetramer is extremely stable, the monomer, once formed, has similar stability (*c*_m_ (refold) = 3.3 M urea) to that of huTTR and is therefore susceptible to denaturation and agitation-induced aggregation. The unfolding kinetics of hbTTR are similar or even slower than those of murine TTR and the protective T119M variant of huTTR, which display hysteresis in their denaturation and refolding profiles and are non-amyloidogenic in acid conditions due to the high kinetic stability of the tetramers (*31, 34*). In contrast to T119M (*31*), the natural species variants studied in this work did not have a kinetic barrier in the re-association of the tetramer during refolding.

The aeTTR tetramer unfolds and refolds reversibly in urea, with a higher *c*_m_ value than huTTR (*c*_m_unfold_ = 4.7 M urea for aeTTR versus 3.3 M urea for huTTR, Table 3). The rate of tetramer dissociation, extrapolated to 0 M urea, is substantially slower for aeTTR than for huTTR (*k* = 0.005 h^-1^ and 0.042 h^-1^, respectively) although it rapidly increases as a function of denaturant concentration (Table 3). The high kinetic stability of the aeTTR tetramer in unperturbed conditions is not able to protect against aggregate formation, since aeTTR aggregates much more rapidly than huTTR, but the high thermodynamic stability of the protomer is consistent with a low propensity to form fibrillar aggregates (Figures S4-S7, Table S3).

Given the large number of substitutions between variants, it is unlikely that the observed differences in aggregation propensity can be reduced to the effect of a single point mutation but arise instead from the network effect of many substitutions. This is manifest in differences in macroscopic properties such as hydrophobicity and overall charge (Table S2). In particular, aeTTR is only weakly charged at pH 7.0, which likely explains its propensity to aggregate rapidly under agitation, possibly without the need for tetramer dissociation. The susceptibility of a given variant towards structural remodeling and fibril formation will depend on the thermodynamic and kinetic stability of the tetramer and the thermodynamic stability of the monomer. These properties will be modulated in complex ways by substitutions that affect hydrophobic packing, hydrogen bonding, and electrostatic interactions, both within the protomer and between subunits of the tetramer. For both hbTTR and aeTTR, there are substitutions in the subunit interfaces that might explain the increased stability of the tetramers against urea denaturation relative to the human TTR. In hbTTR, there are substitutions of huTTR residues H90Y, E92D, and K70R that would potentially stabilize the strong dimer interface and hence the tetramer (Figure S1B). Similar substitutions (H90F, E92D, K70R) are also found in the non-amyloidogenic mouse TTR. Electrostatics calculations in fact indicate that H90 and E92, which are unique to huTTR in our natural variant set (Figure S1C), and to a lesser extent K70, destabilize the strong dimer (*36*). In particular, the carboxyl groups of E90 and E92’, across the strong dimer interface, are only 2.7Å apart (*37*) and are strongly destabilizing; substitution with the shorter D92 side chain in hbTTR and mouse TTR may increase the distance between the carboxylates and hence stabilize the dimer by decreasing electrostatic repulsion. In aeTTR, the substitutions L17M and V121I would be expected to enhance hydrophobic packing across the weak dimer interface and hence stabilize the tetramer, likely contributing to the substantial increase in the midpoint for urea unfolding (c_m_ = 4.7 M urea *versus* 3.3 M urea for unfolding of aeTTR and huTTR, respectively).

The oligomeric aggregates observed in negative stain TEM appear to constitute an intermediate in the aggregation pathway of all TTR variants (Figure 5). They resemble TTR aggregates that form transiently at a very early stage (∼ 9 min) of acid denaturation (*32*). Under the neutral pH conditions in this work, the oligomers appeared to be stable for at least a few hours in quiescent conditions after harvesting samples post-agitation. Of the variants, aeTTR formed spherical particles most rapidly. The particles were comparatively uniform in size (∼15-30 nm diameter) and resembled the mesoscopic clusters observed during aggregation of globular proteins such as p53 (*38, 39*). The p53 clusters are formed by accumulation of misassembled oligomers and act as sites for nucleation of fibrils. The oligomeric TTR clusters appear to be on-pathway in the progression towards large aggregate or fibril formation, since they disappear as aggregation continues.

The variable outcomes of transthyretin aggregation for three physiologically diverse species (human, hummingbird, African elephant) is, to the best of our knowledge, the first experimental indication that serum proteins such as transthyretin may have adapted during evolution to variations in blood flow characteristics. The diverging aggregation kinetics of the variants supported species-specific adaptations of TTR to different heart rates, but the observation of different morphologies of aggregates (amorphous versus fibrillar) demonstrated that multiple steps along the aggregation pathways must be considered for a complete description of the molecular mechanism. In future work, it will be important to extend the studies to transthyretin from additional species with differing heart rates (Figure S1) to consolidate the evidence for evolutionary adaptation. Disentangling the effects of the numerous sequence variations will be challenging since mutations are likely to have co-evolved in networks, with compensatory substitutions that ensure retention of function and stability concordant with the physiological requirements of the organism.

## Materials and Methods

### Preparation of proteins and standard agitation conditions

Recombinant huTTR was overexpressed and purified as previously described, using a Ni-affinity tag protocol described previously (*12*). The genes for hbTTR (Uniprot A0A091I5I6) and aeTTR (Uniprot G3TDX2) were obtained as cDNA from Integrated DNA Technologies and were cloned with overlapping primers into the same His_6_-fusion construct as huTTR. The variant sequences that were used in this work and an amino acid composition analysis can be found in Tables S1 and S2. All TTR concentrations in this work are calculated based on protomer molar UV extinction coefficients (Table S2). Protein identity and purity were confirmed by MALDI-MS, SDS-PAGE and HPLC (Figure S3). Reducing agent (0.5-2 mM TCEP) was used throughout the purification and in the final high-concentration protein stocks (>300 μM TTR) to ensure a fully reduced starting condition, but was not replenished after dilution to the physiological range for agitation experiments. TTR samples were agitated with previously reported conditions (*12*) (1200 rpm, Teflon stir bar, 500 μl sample volume, ambient temperature) in agitation buffer (10 mM potassium phosphate, pH 7.0, 100 mM KCl, 0.5 mM EDTA).

### Native electrophoretic mobility shift assay (native gel)

The starting concentration for all samples in the native gel experiment was *c*_0_ =10 μM. Samples were agitated at room temperature for 90 h. A reference sample was kept on the bench without agitation. After 90 h, samples were centrifuged to remove the insoluble fraction (15 min at 17k x g). Native gel loading dye was added to the supernatant (final ∼10% glycerol, <0.01 % bromophenol blue dye) and 15 μl samples were loaded onto an agarose gel (1% w/v in 25 mM Tris, 192 mM glycine, pH 8.3). Electrophoresis was performed at 80 V for 40 min, followed by staining with Coomassie solution (10% methanol, 50% acetic acid, 0.01% Coomassie blue) and de-staining in deionized water.

### Agitation-induced aggregation assay

The main agitation assay was performed as previously described (*12*). To avoid errors from small variations in protein and buffer batches, comparative TTR variant screens were run in parallel with common stock solutions and *c*_0_ in the range from ∼5 μM - 20 μM. For 96-well plate-based light extinction and fluorescence assays we used UV-transparent, opaque wall plates (Greiner, UV-star, Item No. 655809) for both the ‘total’ analysis and the ‘supernatant’ analysis (30 min at 17k x g to remove large aggregates). Light scattering was measured at 330nm (*A*_330(t)_), and 600 nm (*A*_600(t)_). Supernatant protein concentration was measured by extinction at 280 nm (*A*_280(s)_) on a Tecan Infinite® 200 PRO plate reader (100 μl sample per well). All plate reader data were baseline-corrected using reference data from buffer-only wells.

To compare the aggregation kinetics of the TTR variants, agitation time-course experiments were performed at room temperature and neutral pH following recently established protocols (*12*). Independent samples from common stocks of TTR, for 3 replicates per time point, were agitated by rapid stirring for up to 240 h. At each time point, the agitated samples were split into two fractions for analysis. One aliquot was used as is (total fraction ‘t’) to determine the aggregate content based on light scattering at 330nm (*A*_330(t)_) and 600 nm (*A*_600(t)_) and to measure the fibril content by the increase in thioflavin T fluorescence (ThT gain). The second aliquot was centrifuged to remove large aggregates; the total amount of residual soluble protein in the supernatant (soluble, ‘s’) was determined by UV extinction at 280 nm (*A*_*2*80(s)_), and the remaining TTR tetramer fraction was determined using the resveratrol (RV) binding assay (RV added post-agitation, final 33 μM RV, 10 min incubation in the dark without shaking). The RV concentration was kept constant throughout the time course. Light scattering time course profiles were fitted with Equation S1 as described previously (*12*) to yield the midpoint time *T*_m_, the exponential phase rate *k*, and the plateau intensity *I*_p_. Decaying *A*_280(s)_ time course profiles were fitted with equation S2.

### Thioflavin T assay

After measurement of total absorbance at 330 and 600 nm (*A*_330(t)_ and *A*_600(t)_), the samples were immediately reused for the ThT fluorescence assay (excitation wavelength λ_ex_ = 440 nm, emission wavelength λ_em_ = 480 nm) by addition of ThT (ThT added post-agitation, final 100 μM from 1 mM stock in agitation buffer, 10 min incubation in the dark without shaking). Individual ThT fluorescence intensities were normalized to ThT gain = *I*_ThT_*/I*_ThT,free_, by the buffer-only ThT signal. ThT fluorescence time course profiles were fitted with Equation S1.

### Resveratrol binding assay

After measurement of absorption and light scattering of the ‘supernatant’ fraction, samples were immediately reused to determine the tetramer content by a resveratrol (RV) binding assay (*33*). RV was added to a final concentration of 25 μM (>5-fold excess of RV over initial tetramer concentration at highest *c*_0_) from 100 mM stock in DMSO. RV fluorescence intensity (λ_ex_ =350 nm, λ_em_ =390 nm) was measured after 10 min incubation in the dark. The presence of TTR tetramer was monitored by the increase in fluorescence intensity relative to free RV in agitation buffer (RV gain = *I*_RV_/*I*_RV,free_). Time course profiles were fitted with Equation S2.

### Negative stain transmission electron microscopy (TEM) and mass photometry

Samples (c_0_ =2.5 μM) were agitated for 24 h, as described above, and two independently stirred replicates per variant were pooled for oligomer analysis. Very large aggregates were removed by gentle centrifugation (2 min at 2000 x g). Mass photometry was performed on a Refeyn TwoMP instrument by rapid manual dilution (1:40) to a final protomer concentration of 50 nM (0.5 μl TTR sample added to 20 μl buffer). Additional details can be found in the Supporting Information. The same pooled samples (3 μl) were loaded without dilution onto plasma activated carbon-coated copper grids, incubated for 1 min, blotted, washed with two cycles of 3 μl drops of distilled water, and finally stained with 2% uranyl formate solution (three times 3 μl drops, ∼2 minutes total stain time). Images were acquired with 200kV acceleration on a Talos F200C G1 microscope. Crystallization of stain in the presence of residual phosphate buffer occurred in parts of the grid but adequate areas for automated image acquisition could be found on each grid. Particle statistics were obtained by analysis of the grids using DoG Picker (*40*). Particles of aeTTR in the size range of oligomers (200-500 Å) were classified using cryoSPARC blob picker, which picked ∼54000 particles from 74 micrographs. Particles were further filtered to retain only bright, homogeneous large particles (NCC score >0.48, local power >∼15000), resulting in 2059 extracted particles that were aligned in 50 classes.

### Imaging of aggregates

Samples (c_0_ =10 μM) were agitated for 74 h. For ThT fluorescent imaging, 0.5 μl 1 mM ThT stain was preset in a silicone gasket well (Grace Bio-Labs, GBL103250) on a cleaned coverslip and mixed with 6.5 μl total TTR sample (no centrifugation) by gentle pipetting. Slides were then covered with a coverslip to prevent evaporation and left to sediment for 60 min. Images were acquired at the glass surface using an epifluorescence microscope (Nikon Ti2-E automated inverted microscope) with either a 100x oil immersion objective (CFI60 plan apochromat lens, N.A. 1.45) or with a 40x water immersion objective (CFI60 Apochromat Lambda S LWD, N.A. 1.15), using a Kinetix sCMOS camera (Teledyne Photometrics) and 5.4% 440 nm laser power (60 ms exposure). For unbiased statistics, an automated image stack of ten non-overlapping locations within each well was acquired. Images containing parts of the well edge were removed from the stacks, resulting in collection of images covering a total glass surface area of ∼3.5-4 mm^2^ for the 40x magnification, and ∼0.38 mm^2^ for the 100x magnification. Image stacks were loaded into ImageJ (*41*) and a particle mask was calculated using the ‘Moments’ filter (threshold 5500-64000 for 40x, and 3000-64000 for 100x). Particle statistics were calculated with the ‘Analyze Particles’ function, using a particle area range of 0.05-10000 μm^2^ and allowing all circularities. Enhanced resolution ‘pseudo-confocal’ images were collected on a selected sample (huTTR, c_0_ = 20 μM, 72 h agitation, stained with 100 μM ThT) using a Keyence BZ-X800 microscope (GFP filter cube, 100x oil immersion objective) and Fourier plane filtering (1D filtering).

### Urea denaturation assay

Urea denaturation assays were performed using established protocols(*31, 33, 42*). Briefly, TTR samples (10 μM) were incubated at ambient temperature (∼25 °C) in agitation buffer supplemented with 0.5 mM TCEP and twelve urea concentrations (0-8 M, 3 replicates per condition) in sealed 96-well flat bottom plates (Greiner, UV-star, Item No. 655809). Native Trp fluorescence was measured after 120 h on a BioTek Synergy neo2 multi-mode reader (Agilent) with λ_ex_ = 280 nm, and λ_em_ = 335 nm and λ_em_ = 355 nm. The intensity ratio *R*_355/335_ =*I*_355nm_/*I*_335nm_ was calculated after background subtraction (buffer-only measurement). Denaturation curves were fitted with the models of Hurshmann-Babbes et al. (*33*) (see Supporting Information). The same plates were re-used to monitor tetramer dissociation by RV binding with the protocol described above for the agitation assay.

### Urea refolding assay

TTR was refolded in urea following denaturation in a stronger denaturant according to established protocols (*34*). TTR variants at ∼350-530 μM concentration were denatured in guanidinium hydrochloride (10 mM potassium phosphate, pH 7.0, 6 M Gdn-HCl, 100 mM KCl, 1 mM EDTA) for 48h at ambient temperature in individual dialysis containers (Spectra Por S/P 3 Dialysis Membrane, 3.5 kDa MWCO). Dialysis bags were then transferred to 8M urea buffer and dialyzed for ∼12h to remove Gdn-HCl. TTR recovery after unfolding was > 90% and unfolded samples remained clear, without signs of aggregation. Samples were diluted to 130 μM in 8M urea buffer and further diluted to 10 μM into preset urea buffers at concentrations from 0 M-8M urea on a 96 well plate. Refolding was evaluated by Trp fluorescence intensity on the plate reader after 24 h and 120 h. RV fluorescence intensity gain was determined on the same samples after 120 h.

## Acknowledgments

We thank Euvel Manlapaz for excellent technical assistance and Gerard Kroon for NMR facility management.

## Author contributions

Conceptualization: I.R., P.E.W.

Methodology: I.R., D.S., M.R., M.A.M.-Y.

Investigation: I.R., D.S., M.R.

Visualization: I.R., D.S., M.R.

Supervision: G.C.L., H.J.D., P.E.W.

Writing - original draft: I.R.

Writing - review & editing: H.J.D, P.E.W.

