## Supplementary Material for "From Hummingbird to Elephant: Amyloid Formation in Natural Transthyretin Variants"

Irina Ritsch *et al.*

**This PDF file includes:**

Supplementary Text
Figs. S1 to S15
Tables S1 to S3
Supporting videos S1-S3

**Supplementary Text**

**Additional Experimental Details**

Matrix-assisted laser desorption mass spectrometry (MALDI-MS) and reverse phase high pressure liquid chromatography (HPLC)

TTR stocks (100 μM in agitation buffer) were diluted 1:20 in 0.1% trifluoroacetic acid (TFA) in double distilled water. 1 μl diluted TTR was spotted with 1 μl saturated sinapic acid solution on a stainless steel target plate. MALDI-MS was recorded on a Bruker MALDI-TOF using linear detection in the range of 5-20 kDa. No significant intensity was detected for higher order oligomers at higher mass ranges, confirming that the tetramer dissociates after dilution into 0.1% TFA. The peaks in HPLC are also expected to be the monomeric forms of the variants. For HPLC, TTR solutions (7 μM, diluted from ~500 μM stock in agitation buffer) were prepared in water plus 0.1 % TFA and loaded (100 μl) onto an HPLC column (bioZen 3.6 μm Intact C4). Samples were eluted (1 ml /min flow) with a gradient from 20 % to 60 % acetonitrile in 0.1% TFA. Eluting protein was detected by UV light extinction at 214 nm.

Calibration of analytical size exclusion chromatography

Analytical size-exclusion chromatography (SEC) was performed as described in the main text using a self-packed Superdex 75 10/300 column (Sigma-Aldrich) with dead volume $V_0$ = 7.5ml, eluted with 10 mM KPi, pH 7.0, 100 mM KCl, 1 mM EDTA, 1 mM DTT buffer at a flow rate of 0.5 ml/min. Calibration was performed using the same column and buffer as for the TTR variants. Elution volumes $V_{elution}$ that correspond to tetramer and monomer were identified by comparison to the chromatograms of a standard protein mixture (bovine thyroglobulin (670 kDa), bovine γ-globulin (158 kDa, $V_{elution}$ = 10.8 ml), chicken ovalbumin (44 kDa, $V_{elution}$ =13.3 ml), horse myoglobin (17 kDa, $V_{elution}$ =15.7 ml), and vitamin B12) in the same buffer. The peak elution volumes of the TTR orthologs were 12.5 ml for human and hummingbird TTR, 12.6 ml for elephant TTR. A very minor shift in the aeTTR elution peak may be due to slightly increased sticking to the column material.

Urea denaturation assay for kinetics

The rate of unfolding of huTTR and aeTTR in urea was determined by monitoring changes in the fluorescence intensity at 355 and 335 nm at 7 time points (2-155 h) over a range of urea concentrations from 0 M to 8 M. Fluorescence intensities were measured on a SpectraMax iD5 spectrometer (Molecular Devices) on a single sealed plate per TTR variant. The fitted unfolding rates $k_u$ at urea concentrations above $c_m$ (where monomer unfolding can be considered irreversible) were used for a linear extrapolation of $\log(k_u)$ versus $c_{urea}$ to determine the urea dependence of the dissociation kinetics (slope $m_u$) and dissociation rate at zero denaturant (intercept $\log(k_{u,0})$).

**Additional Data Fitting Details**

Fitting of aggregation time-course data

Fitting of aggregation time courses was performed with a model slightly modified from that described previously (*12*):

Equation S1:

$$I(t) = \frac{I_p}{(1+e^{-k(t-T_m)})} + I_c$$

where $I$(t) is the increase in $A_{330nm(t)}$, $A_{600nm(t)}$ and ThT fluorescence as a function of agitation time $t$, $I_p$ is the plateau intensity, $k$ is the exponential phase rate, $T_m$ is the midpoint time, and the additionally fitted $I_c$ is the constant offset intensity $I(t = 0 \text{ h}) = I_c$. Decaying signals ($A_{280nm(s)}$) were fitted with the inverse trending Equation S2.

Equation S2:

$$I(t) = I_p \left[1 - \frac{1}{(1+e^{-k(t-T_m)})}\right] + I_c$$

Where $I$(t) is the decrease in supernatant $A_{280nm(s)}$ intensity as a function of $t$, $I_c$ is the constant intensity offset, $I_0$ the initial intensity, $k$ is the exponential rate constant and $T_m$ is the mid-point time of signal decay. In this model $I(t \rightarrow \infty) = I_c$. Note that regardless of the fit model, both ThT gain and RV gain fit are expected to have $I_c$~1, which is the expected value for no signal change with respect to the buffer-only fluorescence reference intensity.

Urea denaturation fitting

Endpoint $R_{355/335}$ was fitted to a model previously used for TTR by Hurshman-Babbes et al. (*33*). For simplicity we define the equations with general parameter names and the fit results are referred to as either ‘unfold’ or ‘refold’ in parenthesis depending on which experimental data was fitted. The model is parametrized as a function of denaturant concentration $c_{\text{urea}}$ with Equation S3.

Equation S3:
$$R_{355/335}(c_{\text{denat}}) = \frac{(\alpha_f + \beta_f\ c_{\text{urea}}) + (\alpha_u + \beta_u\ c_{\text{urea}}) e^{m\frac{(c_{\text{urea}} - c_m)}{RT}}}{1 + e^{m\frac{(c_{\text{urea}} - c_m)}{RT}}}$$

where $c_m$ is the mid-point denaturant concentration, $\alpha_f$ is the intercept of the pre-transition slope, $\alpha_u$ is the intercept of the post-transition slope, $\beta_f$ and $\beta_u$ are the pre- and post-transition slopes, respectively, and $m$ is the urea dependence factor of the exponential phase (expressed in units of molar thermal energy units by normalizing with the universal gas constant $R$ and temperature $T$). Note that in this notation, $m$ must be limited to the negative range for convergence of the fit. For resveratrol (RV) binding we fitted an equivalent equation S4.

Equation S4:
$$R_{RV}(c_{\text{denat}}) = \frac{(R_t + r_t\ c_{\text{urea}}) + (R_f + r_f\ c_{\text{urea}})\ e^{m_{RV}(c_{\text{urea}} - c_{m,RV})/RT}}{1 + e^{m_{RV}\ (c_{\text{urea}} - c_{m,RV})/RT}}$$

where the RV gains for free ($R_f$) and tetramer-bound ($R_t$) RV replace the pre- and post-transition ratios, $r_t$ and $r_f$ replace the pre- and post-transition slopes, $m_{RV}$ is the urea dependence factor and $c_{m,RV}$ transition midpoint urea concentration of tetramer dissociation.

For fitting the time-courses we use a single exponential fit model, Equation S5.

Equation S5 :
$$R_{355/335}(t) = R_{unfolded}\ (1 - \Delta R\ e^{-k_u\, t}),$$

with the endpoint ratio $R_{unfolded}$, the ratio change $\Delta R$, and the unfolding rate constant $k_u$, which was fitted individually to each urea condition. Where sufficient datapoints above 1.1 x $c_m$ were available (aeTTR, huTTR), we extrapolated the fitted unfolding rates to the non-denaturing range using Equation S6.

Equation S6 $k_u(c_{urea}) = k_{u,0}\, e^{-m_u\, c_{urea}}$ ,

with the base rate at 0 M urea ($k_{u,0}$), and the urea dependence $m_u$.

Scattering correction of A280s

Light scattering at 330 nm and 600 nm in the supernatant was used to correct the light extinction measurement at 280nm that were used to back-calculate the residual supernatant TTR concentration $c_s$. We applied the two point estimation using a slightly modified equation that accounted for the different wavelengths in our measurements compared to the published correction protocol.

m=(log A330s-log A600s)/(log(330)-log(600));

b=log A600s-m*log(330);

C=exp(log(280)*m +b);

A280s=A280s(uncorrected)-C;

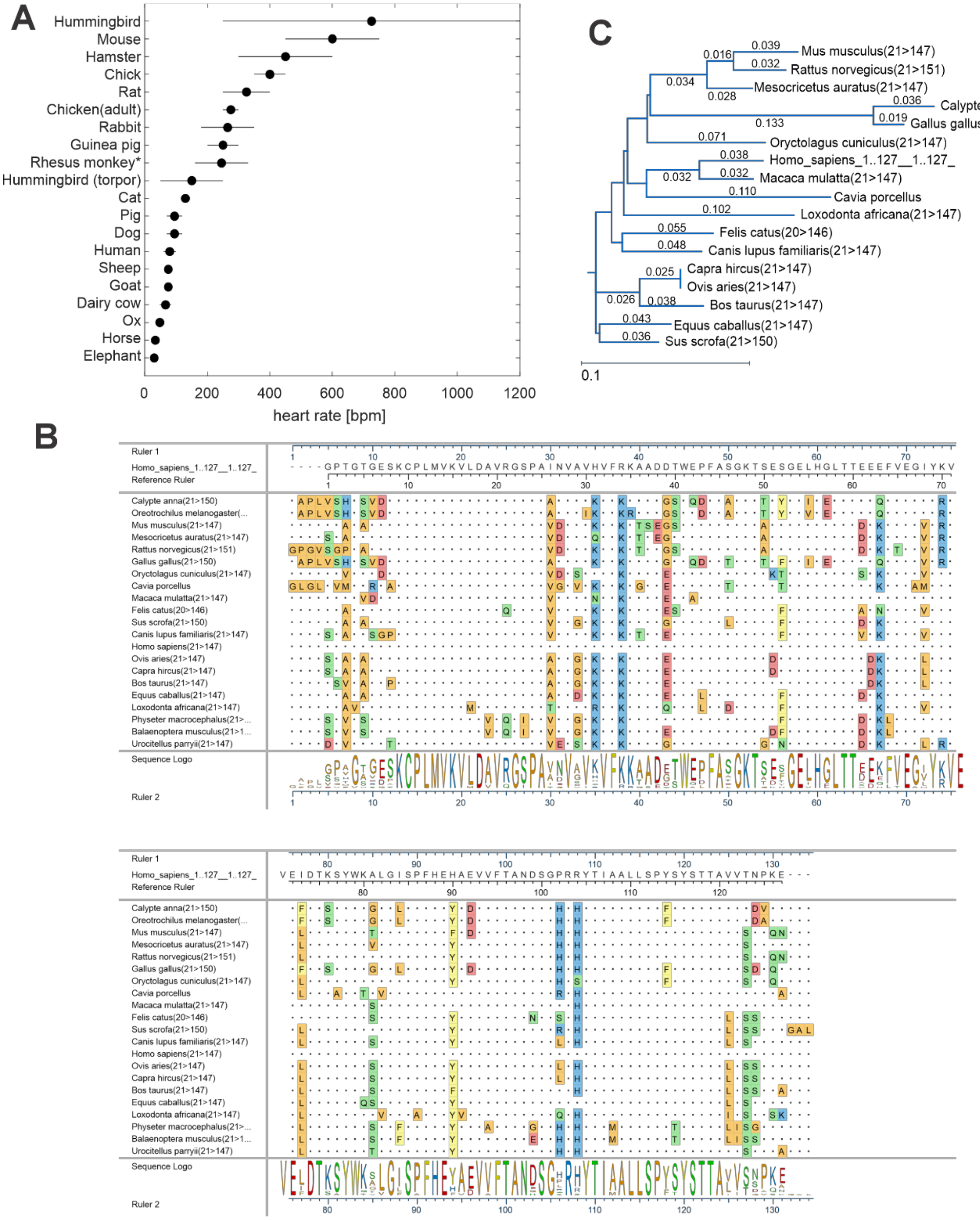


**Figure S1.** A. Reported heart rate ranges across a range of species (*21*). * Rhesus monkey (anesthetized). B. Alignment (clustal Omega) of TTR sequences from species with heart rates reported in panel A. The sequences were downloaded from Uniprot (https://www.uniprot.org/) by searching for “transthyretin” and applying the taxonomy filter. Signal peptides (residues 1-20) were removed prior to alignment. C. Phylogenetic tree of TTR sequences in panel A, generated with BIONJ in DNASTAR.

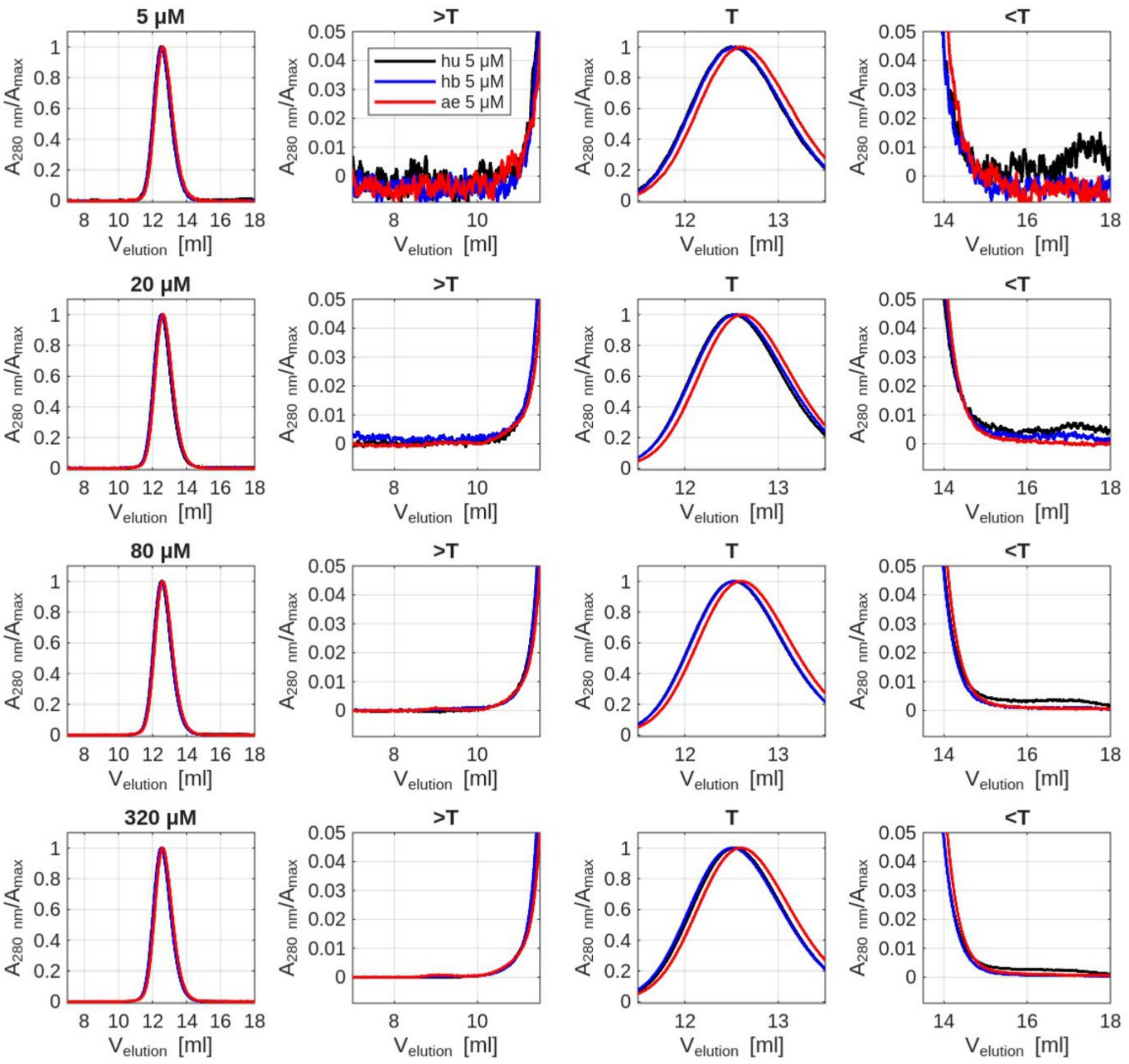


**Figure S2**. Size-exclusion chromatography (SEC) overlays with TTR variants (huTTR: black, hbTTR: blue, aeTTR: red) at four different loading concentrations (rows top to bottom: 5, 10, 80, 320 μM). Dilution on the column is ~1:20 of loading concentration, resulting in on-column concentrations from ~0.25-16 μM. The SEC traces are normalized to maximum intensity, i.e. the tetramer (T) peak intensity. Left to right: full intensity range, zoom on larger than tetramer range (>T), tetramer range (T), and smaller than tetramer range (<T).

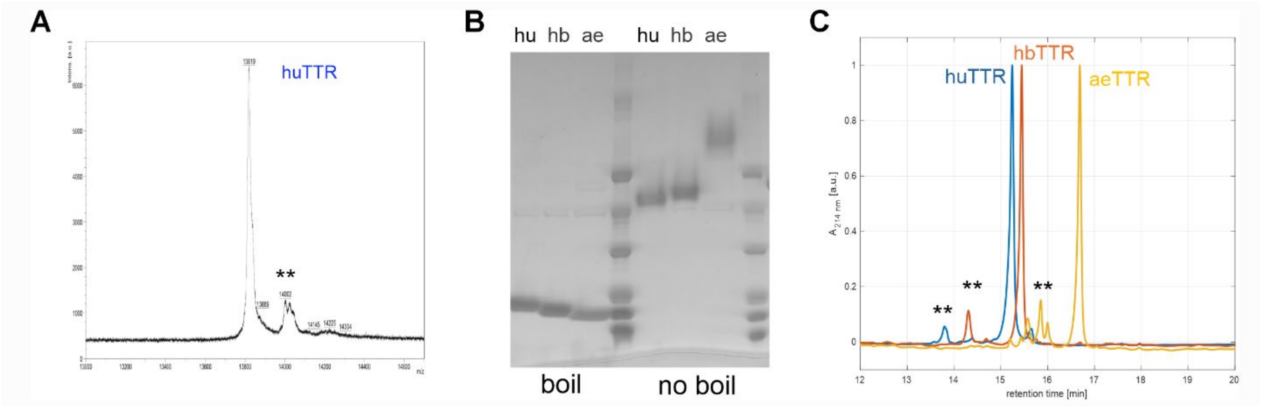


**Figure S3** A. MALDI mass spectrum for huTTR; the double asterisk marks a minor signal of unknown origin at +183 m/z. B. SDS-PAGE gel of TTR species variants. Samples on the left side of gel were boiled 5 min SDS buffer, samples in the right lanes were incubated in the same SDS buffer but were not boiled. C. reverse phase HPLC of purified TTR variants (loaded at 5 µM). The double asterisks mark the position of minor peaks that possibly arise from partial oxidation of Cys or Met.

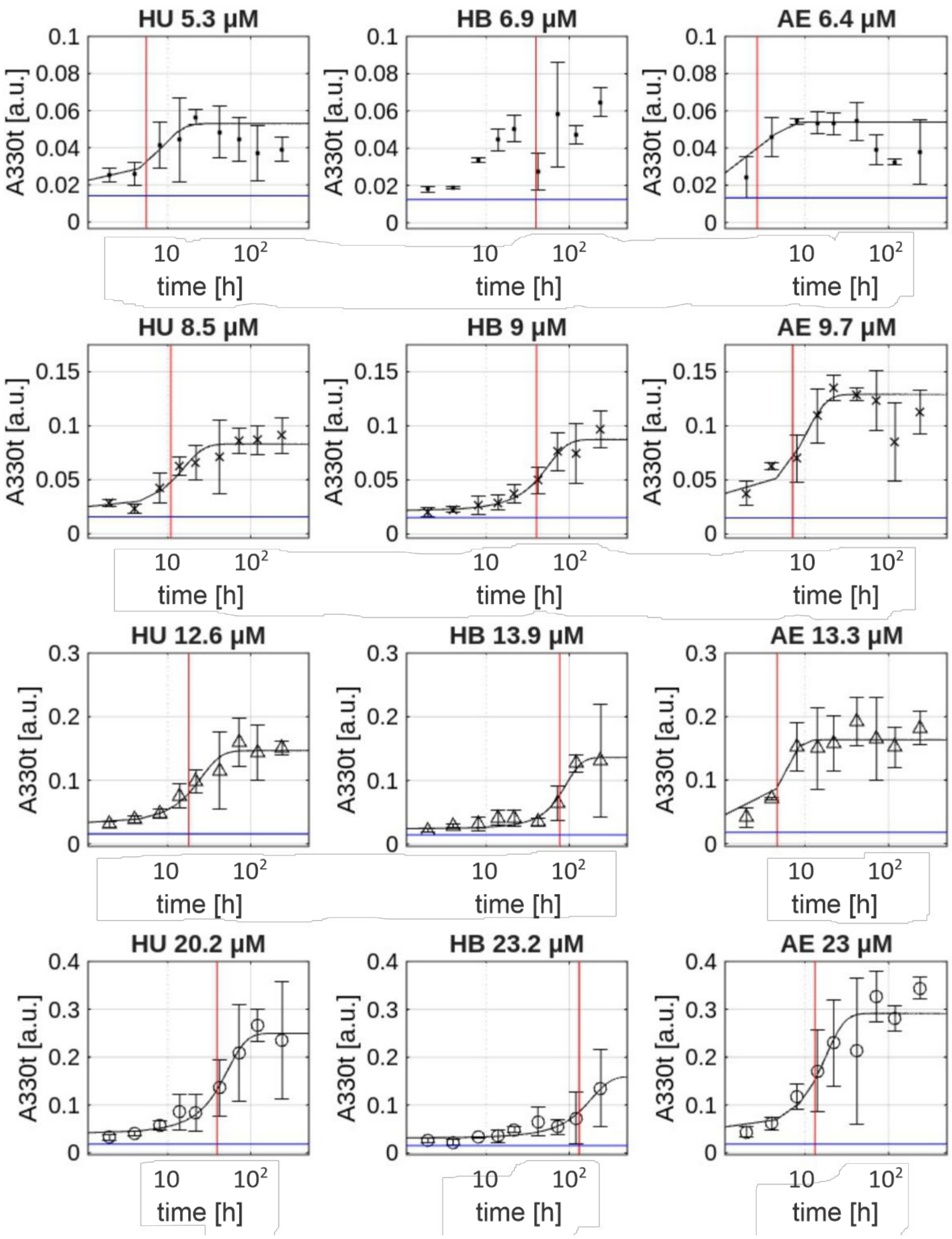


**Figure S4**. Time courses and sigmoidal fits (solid black line) for accumulation of insoluble aggregates of huTTR, hbTTR, and aeTTR measured by light scattering at 330 nm ($A_{330(t)}$) for four different $c_0$ (indicated above plots). The time axis is displayed with logarithmic spacing. The vertical red lines mark the fitted $T_m$ values. the horizontal blue lines mark the experimental $A_{330(t)}$ value at t = 0 h, which cannot be directly displayed in the logarithmic time axis. Data for $c_0$ ~6 µM were not fitted due to insufficient signal to noise.

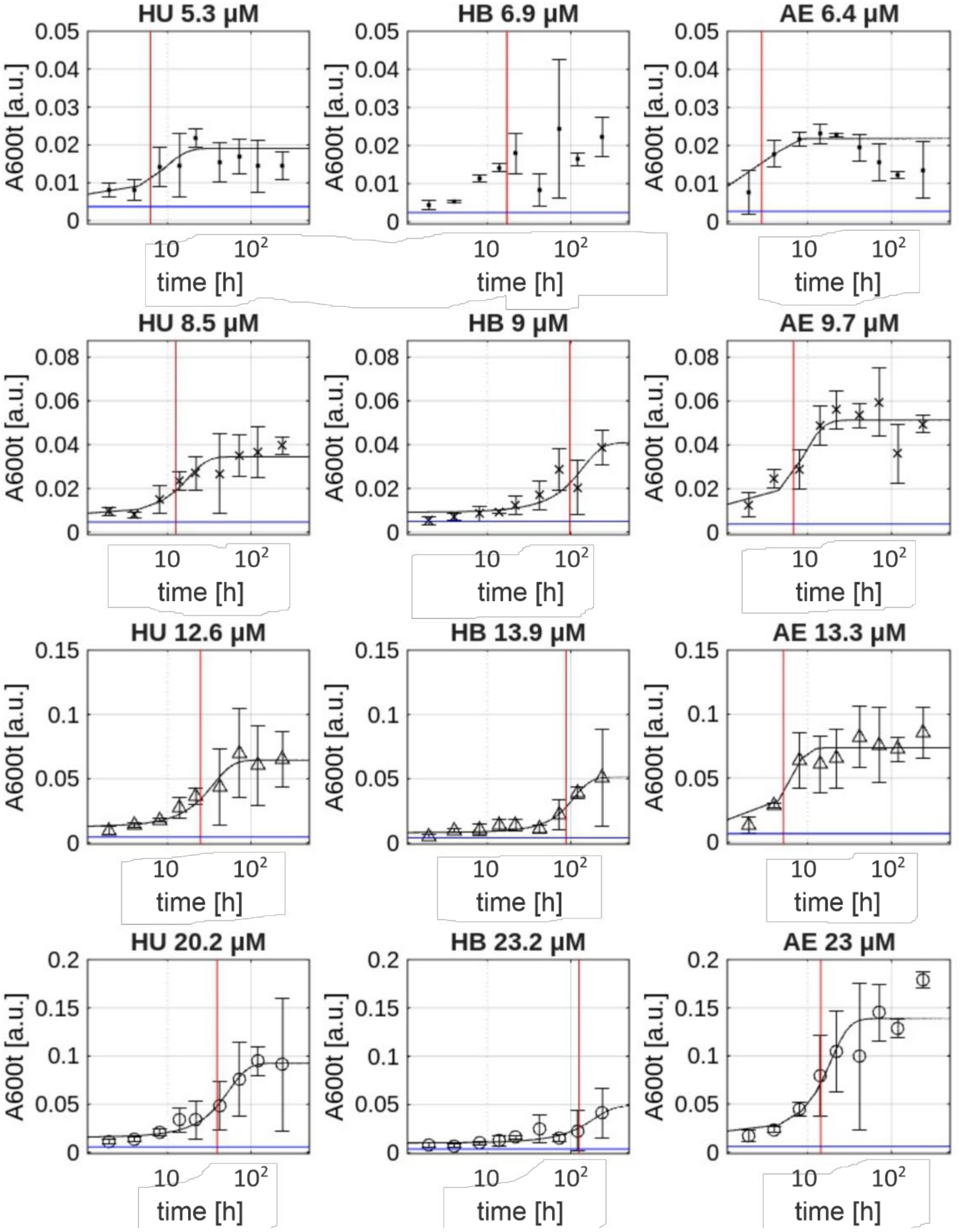


**Figure S5.** Time courses and sigmoidal fits (solid black line) for accumulation of insoluble aggregates of huTTR, hbTTR, and aeTTR measured by light scattering at 600 nm ($A_{600(t)}$) for four different $c_0$ (indicated above plot). The vertical red lines mark the fitted $T_m$ values. The horizontal blue lines mark the experimental $A_{600(t)}$ value at t = 0 h. Data for $c_0 \sim 6$ µM were not fitted due to insufficient signal to noise.

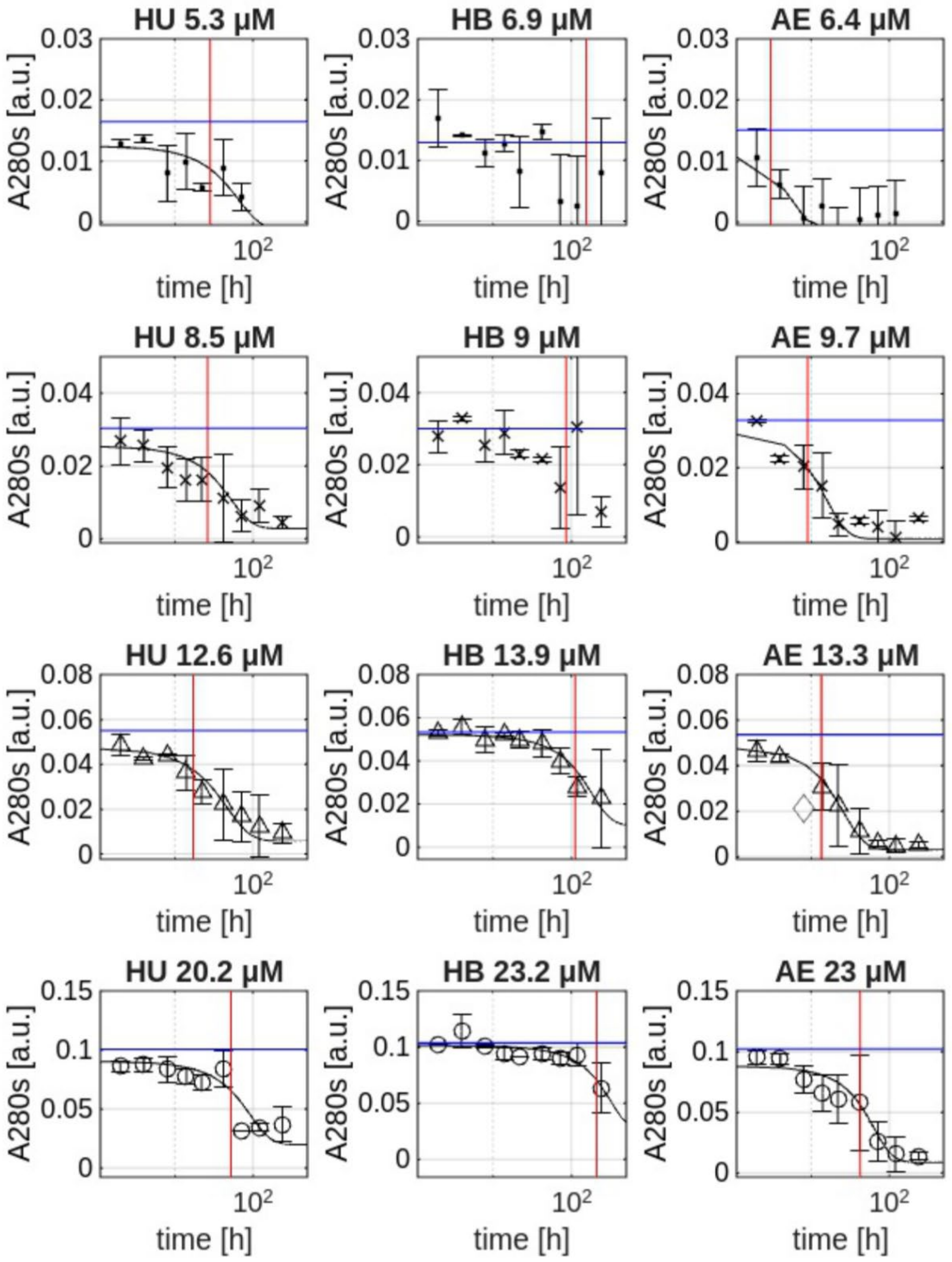


**Figure S6.** Time courses and sigmoidal fits (solid black line) for loss of supernatant of huTTR, hbTTR, and aeTTR measured by absorbance at 280 nm ($A_{280(s)}$) after subtraction of a minor scattering contribution from residual aggregates which was extrapolated from scattering at 330 nm ($A_{330(s)}$) and 600 nm ($A_{600(s)}$) for four different $c_0$ (indicated above plot). The vertical red lines mark the fitted $T_m$ values. The horizontal blue lines mark the experimental $A_{280(s)}$ value at t = 0 h. Fits for data at $c_0$ ~6 μM were not fitted due to insufficient signal to noise. One datapoint in aeTTR was an outlier and omitted from the fit (diamond).

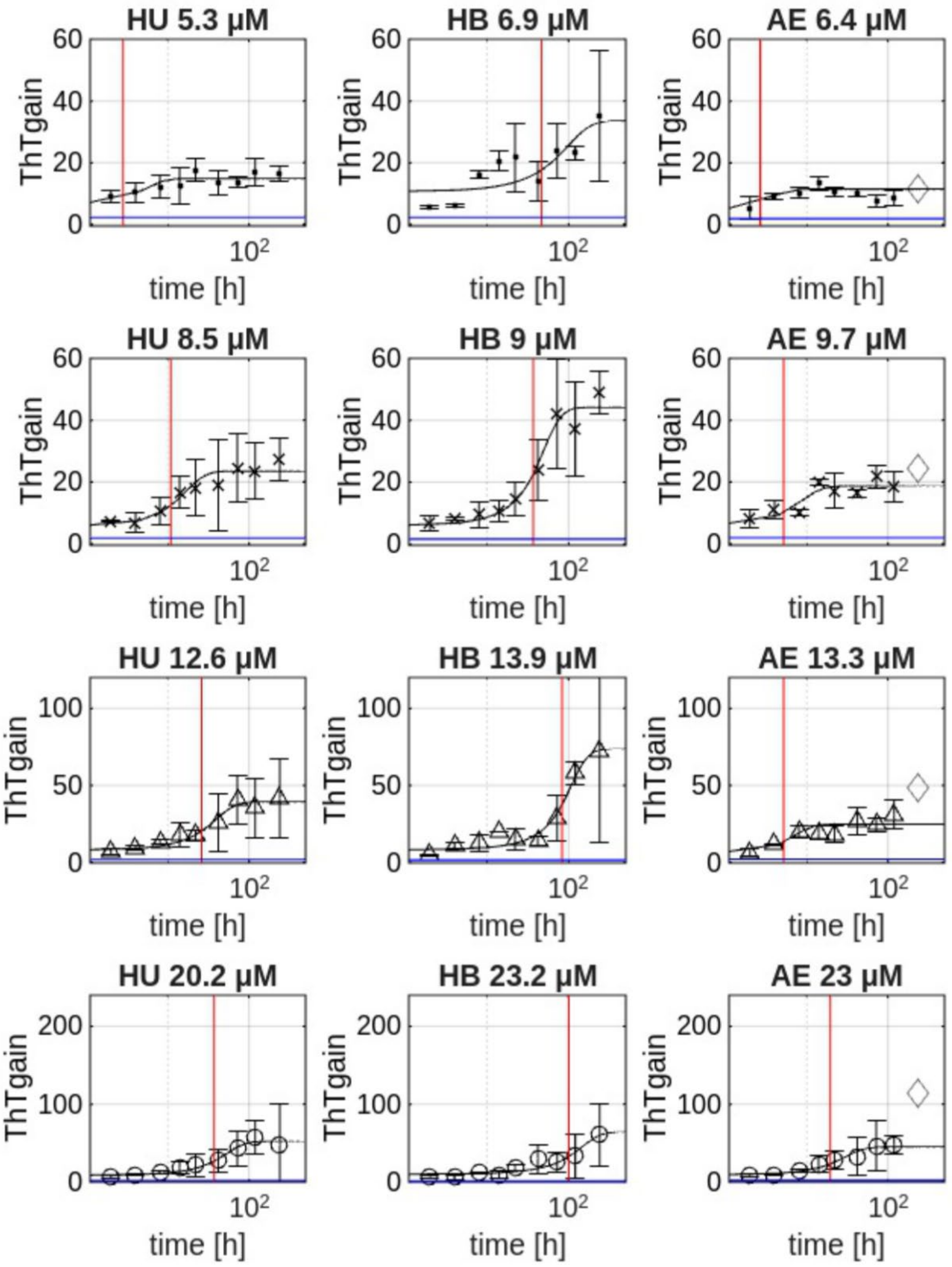


**Figure S7.** Time courses and sigmoidal fits (solid black line) for accumulation of fibrillar aggregates of huTTR, hbTTR, and aeTTR measured by ThT fluorescence intensity gain for four different $c_0$ (indicated above plot). The vertical red lines mark the fitted $T_m$ values. The horizontal blue lines mark the experimental ThT gain at t = 0 h. For aeTTR we omitted the last datapoint (t = 240h, shown as grey diamond) from the fit to model only up to the first plateau of the biphasic behavior.

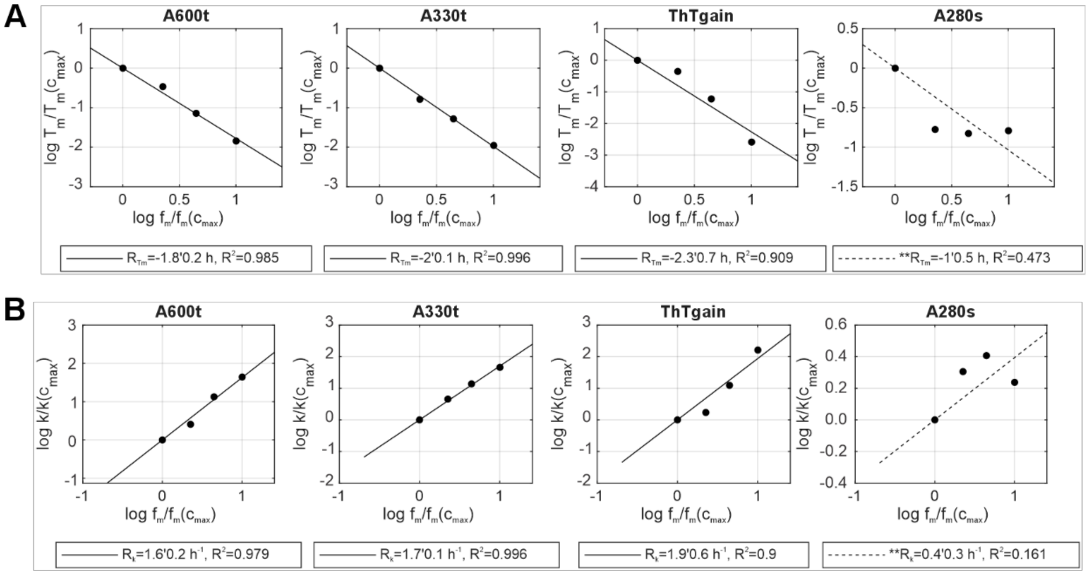


**Figure S8.** Log-log plots of sigmoidal fit parameters versus estimated fraction of monomer $f_m$ for huTTR calculated from tetramer $K_D$ and $c_0$ (*12, 29*).

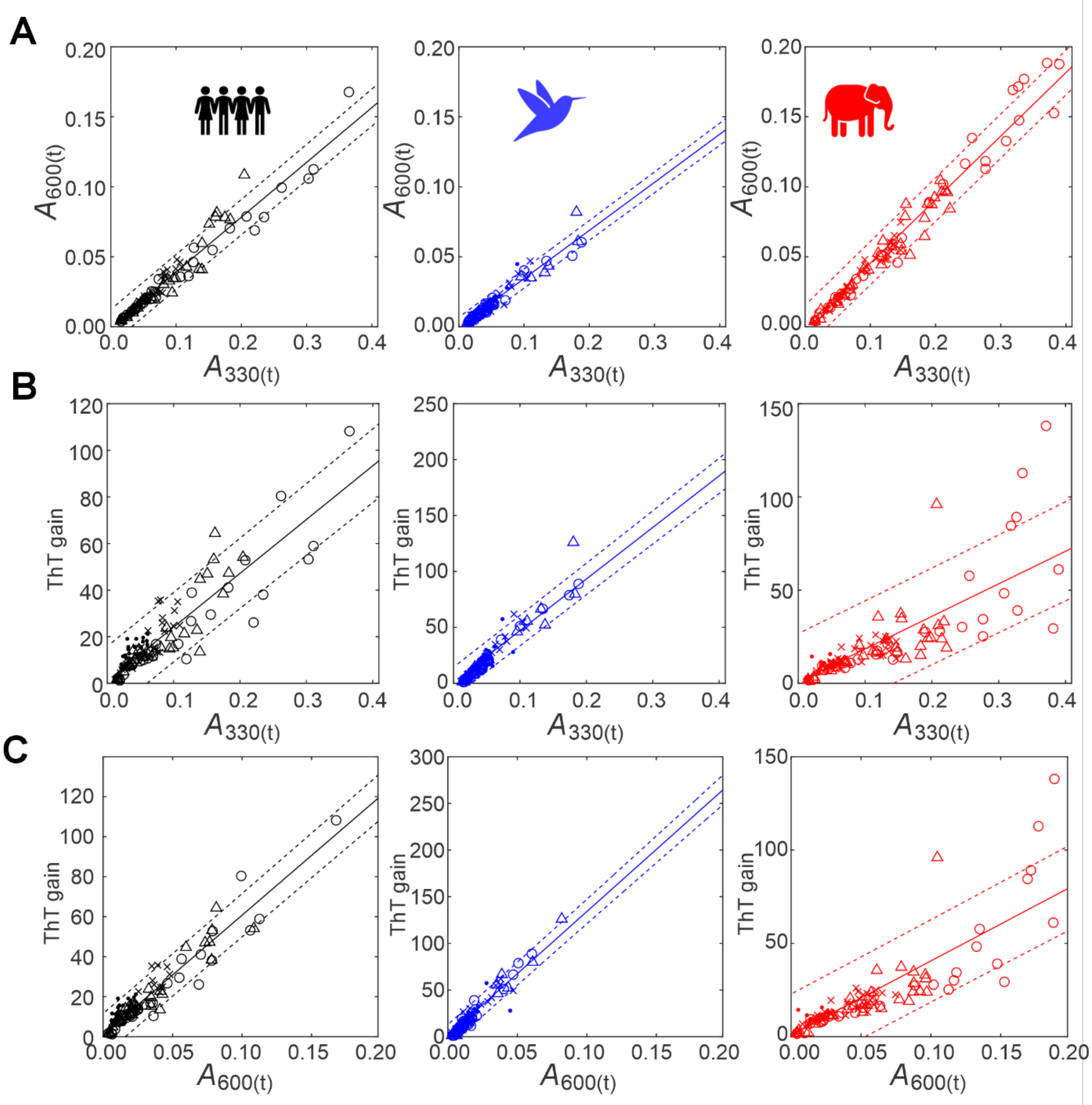


**Figure S9.** A. Correlation between sample scattering at two different wavelengths ($A_{330(t)}$ vs $A_{600(t)}$). B, C. Correlation between ThT fluorescence intensity gain and scattering at 330 nm and 600 nm. Data are plotted for each individually evaluated replicate at all time points and all concentrations ($c_0$: ~5 µM (solid dots),~9 µM (x), ~12 µM (triangles), ~20 µM (open circles)*)* in the kinetic aggregation profiles of Figures S3 S4, and S6 The linear regression slopes and $R^2$ are listed in main text Table 1, only the slope was fitted, offsets were fixed at the theoretical limits, i.e. ThT gain =1, $A_{330(t)}$= $A_{600(t)}$=0, dotted lines show 95% confidence intervals.

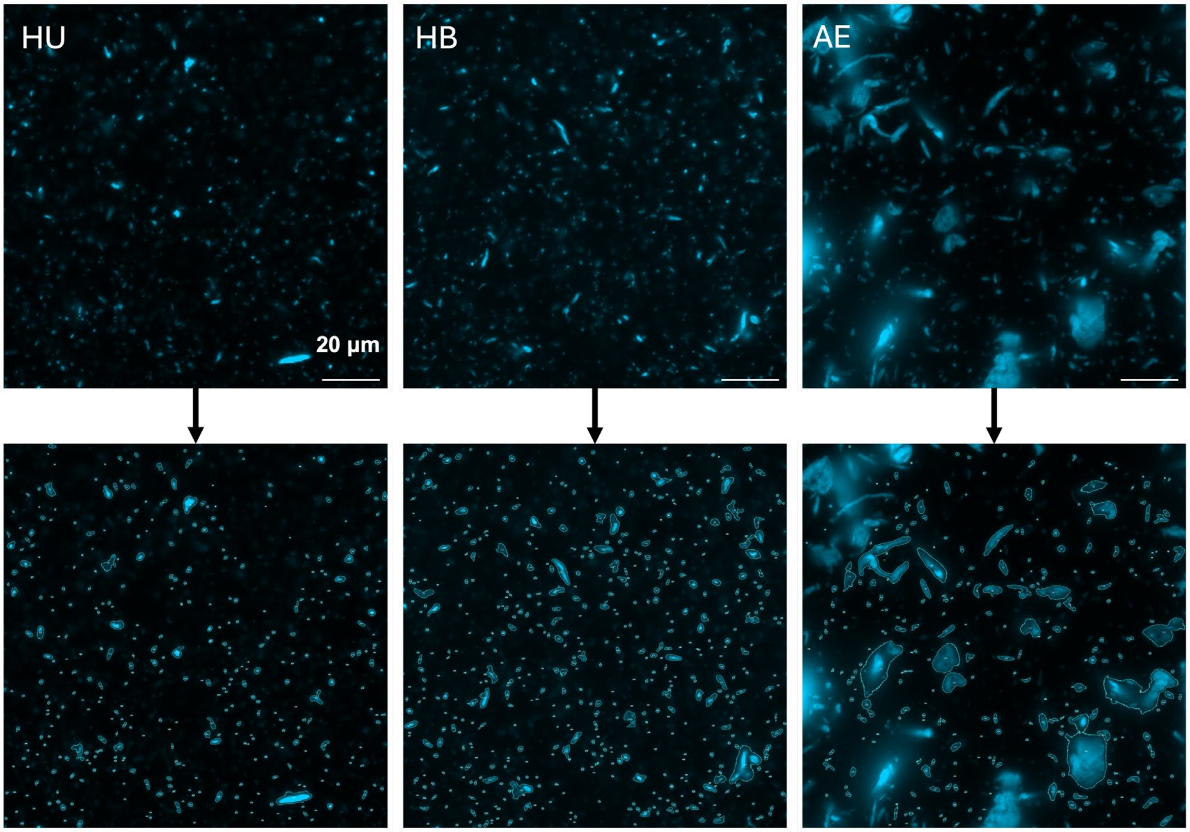


**Figure S10.** Representative 100x magnification ThT fluorescence microscope images of sedimented aggregates formed after 74 h agitation. Particle outlines and particle numbers after thresholding (moments, fixed threshold 3000) are indicated; outlines and numbers may appear faint in print, but can be enlarged in digital version.

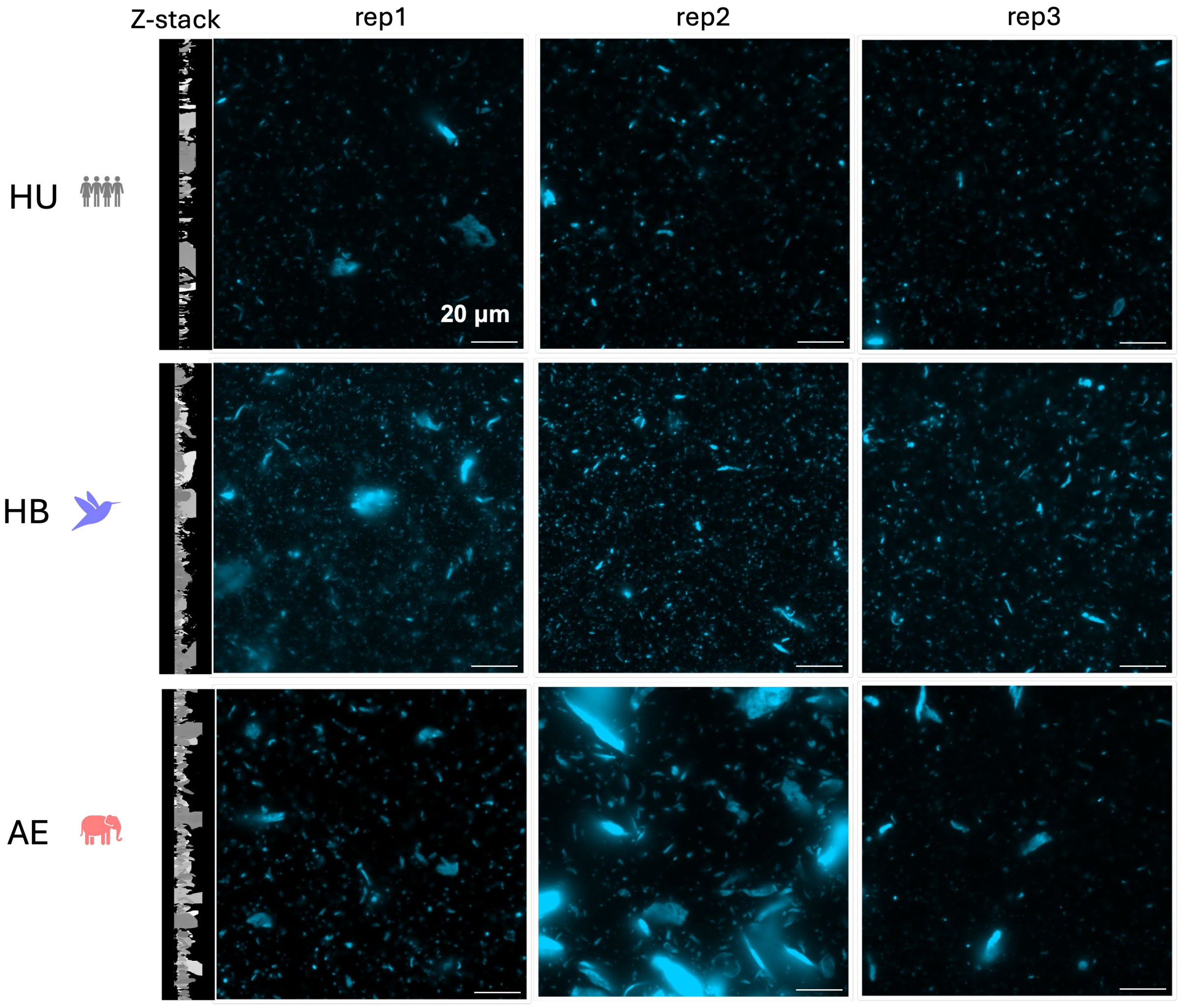


**Figure S11.** ThT fluorescence microscope images of the same samples as in Figure S10 at 100x magnification. Images for replicate 1 were extracted from a z-stack, and the projection perpendicular to the glass surface is shown. In some cases, thin, fibrillar aggregates have a preference to remain oriented perpendicular to the glass surface, which may be partially an artifact of the imaging method.

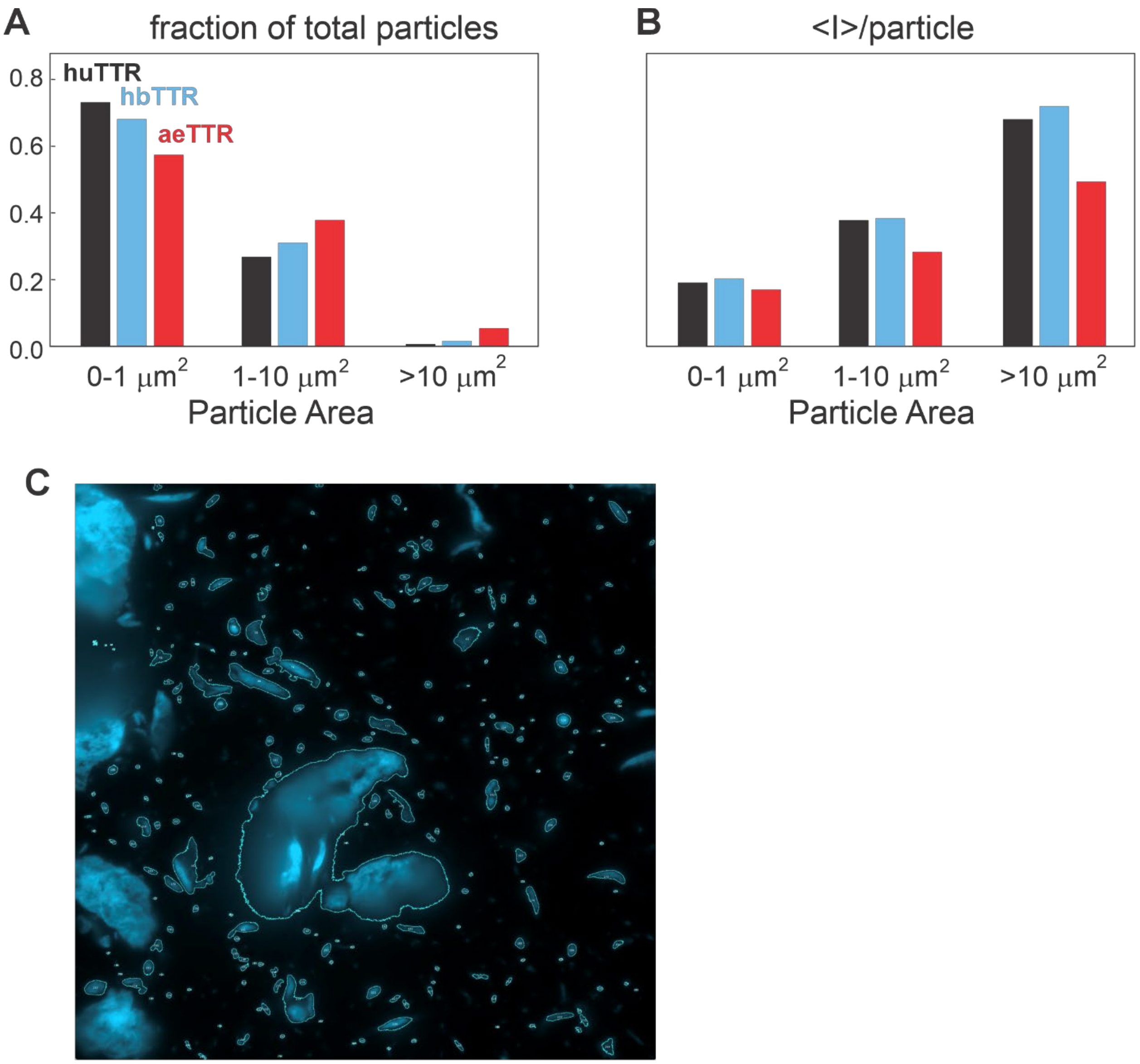


**Figure S12**. Particle analysis of image stacks at 100x magnification, total imaged glass cover slip area was ~0.338 mm$^2$. A. histogram of fraction of particles with particle areas < 1 μM$^2$, 1-10 μM$^2$, and > 10 μM$^2$. B. histogram of average ThT fluorescence intensity per particle *versus* particle area. C. 100x fluorescence microscope image of a large aeTTR particle showing presence of both β-rich and amorphous material that stains only weakly with ThT.

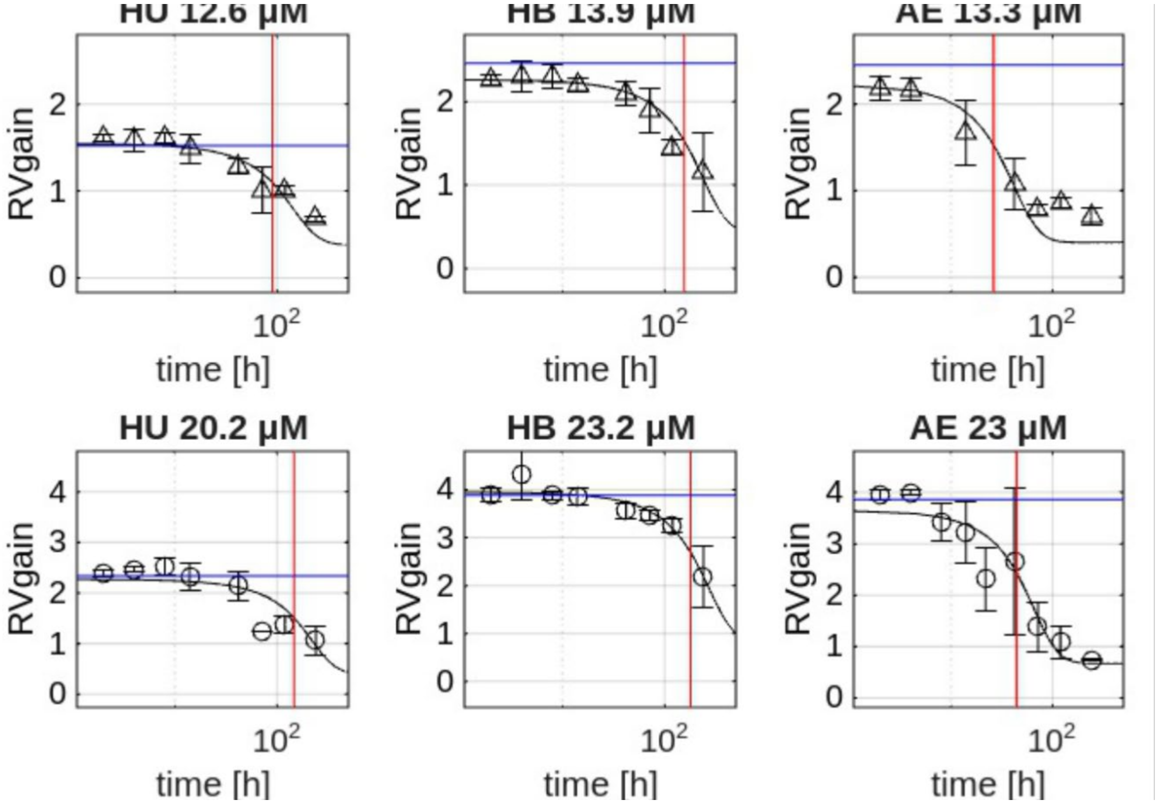


**Figure S13.** Time courses and sigmoidal fits (solid black lines) for loss of supernatant TTR tetramer monitored by RV fluorescence gain for two different $c_0$ (indicated above plot). Plots are shown for huTTR, hbTTR, and aeTTR. The time axes are displayed with logarithmic spacing. The vertical red lines mark fitted $T_m$, horizontal blue lines mark the experimental value at t = 0h, which cannot be directly displayed in the logarithmic time axis. Data for $c_0 < \sim 9$ µM were not fitted due to insufficient signal to noise.

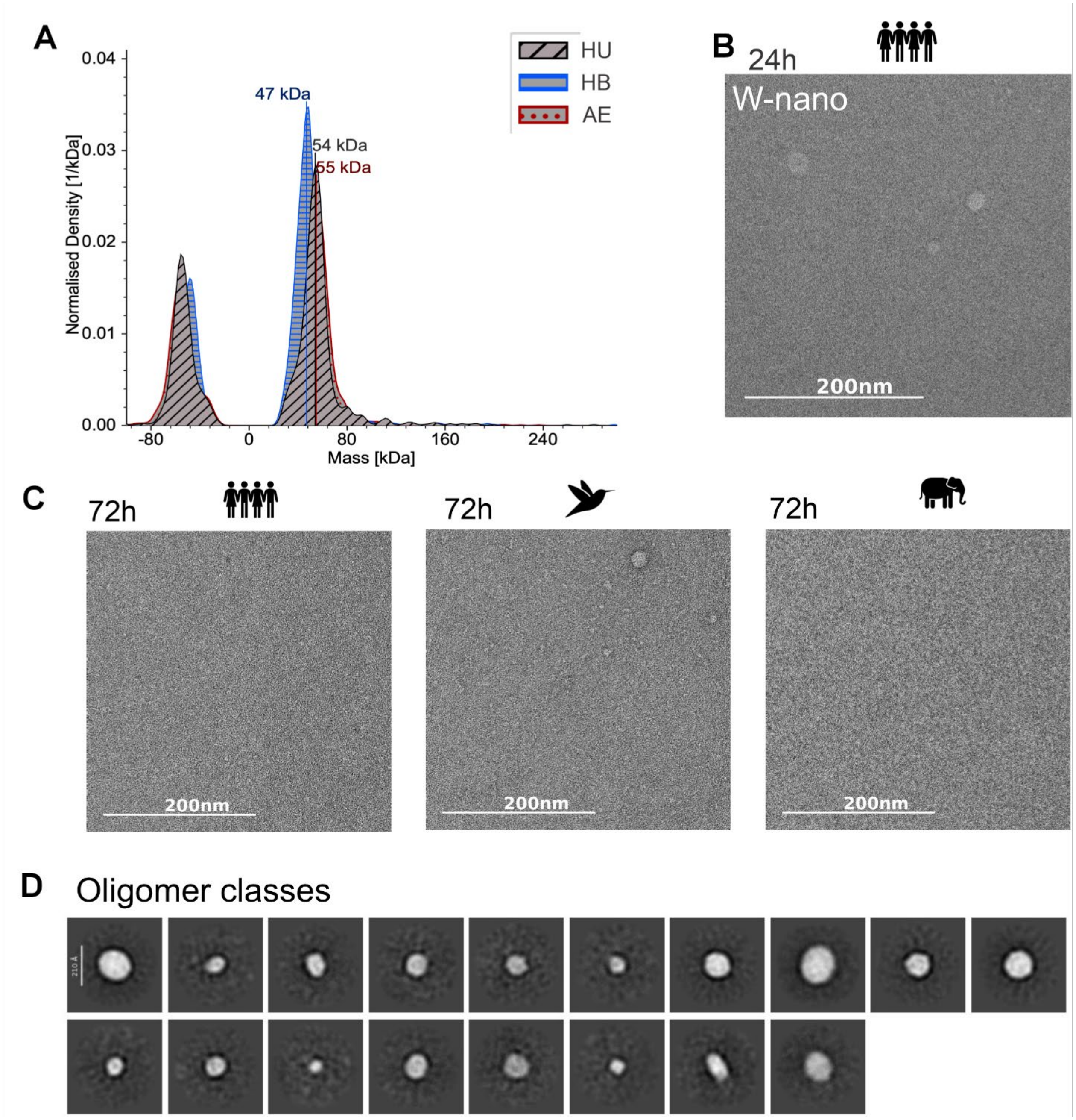


**Figure S14.** A. Reference (non-agitated) mass photometry showing both the binding and unbinding event region, reference samples were diluted 1:50 to avoid surface overloading. B. W-nano stained control for TEM of oligomers (huTTR) after 24 h agitation. C. Uranyl formate-stained TEM of samples after agitation was continued for an additional 48h (total 72 h) showing further disappearance of the tetramers as well as loss of oligomers. D. Additional 2D TEM classes with aeTTR after 24h agitation (not in a particular order).

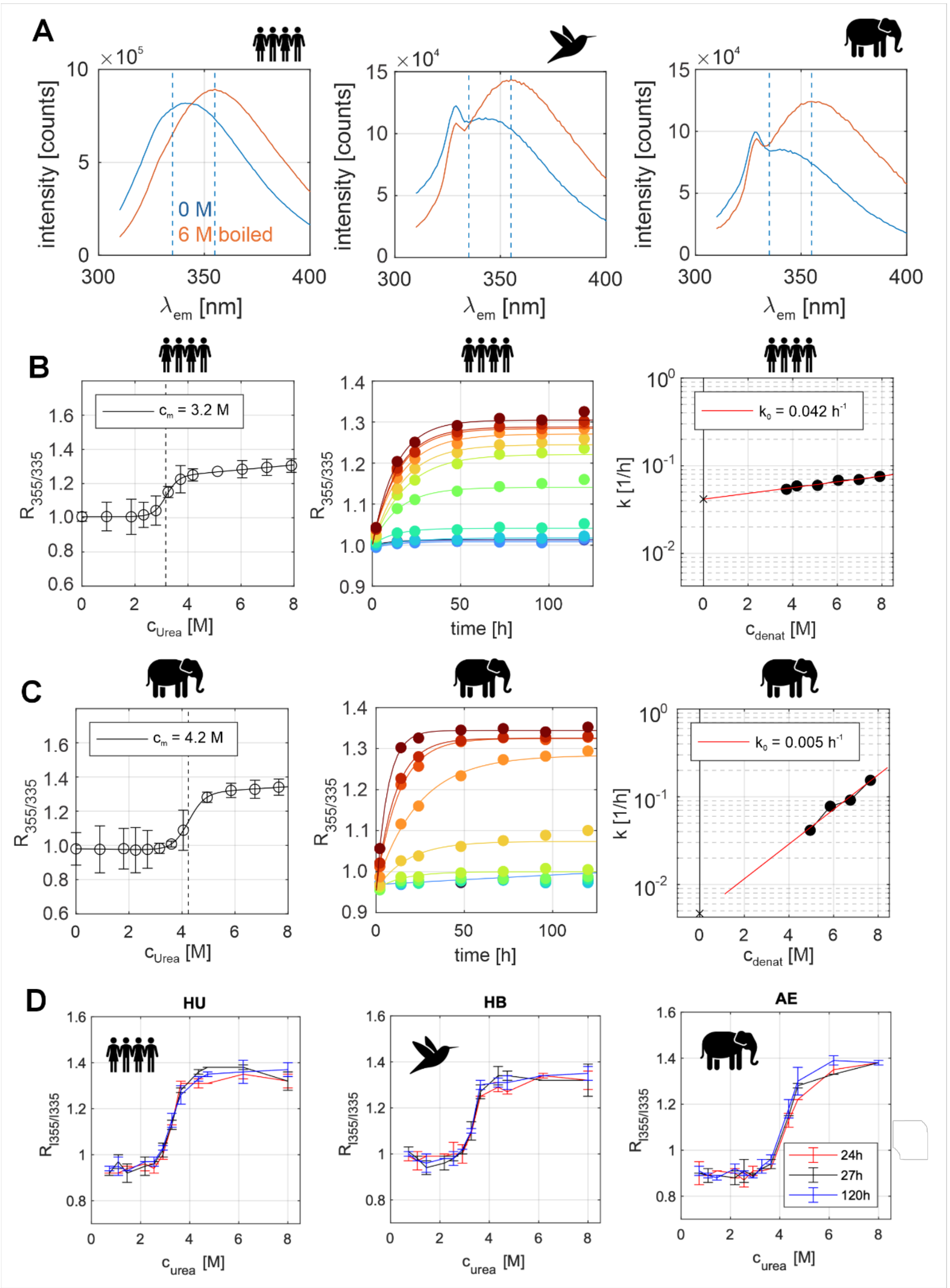


**Figure S15.** A. Tryptophan fluorescence emission spectra of TTR variants. Samples were 1 $\mu$M in agitation buffer (blue) or were boiled for 10 minutes in agitation buffer plus 6 M urea (no turbidity was observed). The vertical dotted lines mark the emission spectra maxima, which are shifted for all TTR variants between the folded and unfolded states. B,C. Urea denaturation of 10 $\mu$M samples of B. huTTR and C. aeTTR. Left panel: changes in R355/335 fluorescence intensity ratio after incubation for 120h in urea at the indicated concentrations. The solid lines show fits to Equation S3 and the dashed vertical lines mark the midpoint $c_m$. Middle panel: kinetics of tetramer dissociation and unfolding at different urea concentrations (blue: 0 M to red ~ 8M) monitored by fluorescence intensity ratio $R_{355/335}$. The solid lines show fits to Equation S5. Right panel: semi-log plot of fitted single-exponential tetramer dissociation rate constant $k$ as a function of urea concentration greater than $c_m$. The slope of a linear fit gives the urea sensitivity $m_u$ and the intercept gives the extrapolated rate $k_0$ at 0M urea (Equation S6). D. Refolding of guanidinium hydrochloride denatured TTR monitored by $R_{355/335}$ recorded after 24h (red), 27h (black), or 120h (blue) incubation in the urea buffer solutions.

**Table S1.**

| **Species common name (scientific name)** | **Short name** | **Entry ID (Uniprot)** | **sequence** |
|---|---|---|---|
| Human (*Homo sapiens*) | huTTR | P02766 | ($G_{-1}$) GPTGTGESKCPLMVKVLDAVRGSPAINVAVHVFRKAADDTWEPFASGKTSESGELHGLTTEEEFVEGIYKVEIDTKSYWKALGISPFHEHAEVVFTANDSGPRRYTIAALLSPYSYSTTAVVTNPKE |
| Anna's hummingbird (*Calypte anna*) | hbTTR | A0A091I5I6 | ($G_{-1}$) APLVSHGSVDSKCPLMVKVLDAVRGSPAANVAVKVFKKAADGSWQDFAAGKTTEYGEIHELTTEEQFVEGIYRVEFDTSSYWKGLGLSPFHEYADVVFTANDSGHRHYTIAALLSPFSYSTTAVVTDVKE |
| African elephant (*Loxodonta africana*) | aeTTR | G3TDX2 | ($G_{-1}$) GPAVTGESKCPLMVKVMDAVRGSPATNVAVRVFKKAADQTWELFADGKTSEFGELHGLTTEEKFVEGVYKVELDTKSYWKAVGISAFHEYVEVVFTANDSGQRHYTIAALLSPYSYSTTAIVSNPSK |

**Table S1**. TTR homolog sequences and sources; ($G_{-1}$) in the sequence column is not part of the natural sequences and represents the additional N-terminal glycine that was introduced for efficient enzymatic cleavage (TEV protease) of the His-tag for purification; this is the equivalent position to $M_{-1}$ in tag-free purifications.

| Short name | # AA | Identity [%] | ϵ [$M^{-1}cm^{-1}$] | MW [Da] | pI | Charge at pH 7.0 | Aliphatic index | GRAVY score | # positively charged (K+R) | # negatively charged (D+E) | # His |
|---|---|---|---|---|---|---|---|---|---|---|---|
| huTTR | 127 | 100 (def) | 18450 | 13818 | 5.31 | -4.8 | 73.70 | -0.274 | 12 | 17 | 4 |
| hbTTR | 130 | 75.59 | 19940 | 14220 | 5.24 | -5.7 | 77.23 | -0.122 | 11 | 17 | 5 |
| aeTTR | 127 | 81.89 | 19940 | 13914 | 5.91 | -2.0 | 75.20 | -0.159 | 13 | 15 | 3 |

**Table S2.** TTR variant characteristics;  for % identity we used the sequences as shown in Table S1 (no signal peptides) and compared to huTTR using blastp (https://blast.ncbi.nlm.nih.gov); the molar extinction coefficient at 280 nm ϵ, the (protomer) molecular weight MW (which includes $G_{-1}$), the isoelectric point pI, the aliphatic index, the GRAVY score and the amino acid statistics were determined by protparam (https://web.expasy.org/protparam/), the charge at pH=7 by protpi (https://www.protpi.ch/).

**Table S3**

| Supernatant fraction measurements | | | | | | | | | |
|---|---|---|---|---|---|---|---|---|---|
| exp | frac | var | $c_0$ [µM] | $k$ [1/h] | $I_p$ | $T_m$ [h] | $I_c$ | $R^2_{adj}$ | Fit eqn |
| A330t | t | HU | 5.3 | 0.3 ± 0.2 | 0.04 ± 0.01 | 6 ± 2 | 0.011 ± 0.004 | 0.855 | S1 |
| A330t | t | HU | 8.5 | 0.2± 0.2 | 0.07 ± 0.02 | 11 ± 4 | 0.012 ± 0.004 | 0.916 | S1 |
| A330t | t | HU | 12.6 | 0.10 ± 0.04 | 0.14 ± 0.01 | 18 ± 5 | 0.012 ± 0.004 | 0.950 | S1 |
| A330t | t | HU | 20.2 | 0.05 ± 0.04 | 0.24 ± 0.06 | 40 ± 10 | 0.013 ± 0.004 | 0.963 | S1 |
| A330t | t | HB | 9 | 0.05 ± 0.05 | 0.08 ± 0.02 | 40 ± 10 | 0.011 ± 0.004 | 0.943 | S1 |
| A330t | t | HB | 13.9 | 0.04 ± 0.01 | 0.12 ± 0.02 | 80 ± 20 | 0.018 ± 0.004 | 0.947 | S1 |
| A330t | t | HB | 23.2 | 0.01 ± 0.01 | 0.15 ± 0.03 | 130 ± 50 | 0.012 ± 0.004 | 0.882 | S1 |
| A330t | t | AE | 6.4 | 1.0 ± 0.2 | 0.044 ± 0.005 | 3 ± 1 | 0.010 ± 0.003 | 0.994 | S1 |
| A330t | t | AE | 9.7 | 0.3 ± 0.1 | 0.12 ± 0.01 | 7 ± 1 | 0.011 ± 0.004 | 0.946 | S1 |
| A330t | t | AE | 13.3 | 0.7 ± 0.5 | 0.15 ± 0.03 | 5 ± 1 | 0.014 ± 0.004 | 0.937 | S1 |
| A330t | t | AE | 23 | 0.2 ± 0.1 | 0.28 ± 0.03 | 13 ± 3 | 0.013 ± 0.004 | 0.886 | S1 |
| exp | frac | var | $c_0$ [µM] | $k$ [1/h] | $I_p$ | $T_m$ [h] | $I_c$ | $R^2_{adj}$ | Fit eqn |
| A600t | t | HU | 5.3 | 0.3 ± 0.5 | 0.02 ± 0.01 | 6 ± 3 | 0.003 ± 0.001 | 0.697 | S1 |
| A600t | t | HU | 8.5 | 0.2 ± 0.2 | 0.03 ± 0.01 | 13 ± 5 | 0.004 ± 0.001 | 0.886 | S1 |
| A600t | t | HU | 12.6 | 0.08 ± 0.03 | 0.06 ± 0.01 | 30 ± 10 | 0.003 ± 0.001 | 0.941 | S1 |
| A600t | t | HU | 20.2 | 0.05 ± 0.04 | 0.09 ± 0.02 | 40 ± 10 | 0.004 ± 0.001 | 0.960 | S1 |
| A600t | t | HB | 9 | 0.02 ± 0.04 | 0.04 ± 0.01 | 100 ± 50 | 0.004 ± 0.001 | 0.736 | S1 |
| A600t | t | HB | 13.9 | 0.03 ± 0.02 | 0.05 ± 0.01 | 90 ± 20 | 0.005 ± 0.001 | 0.951 | S1 |
| A600t | t | HB | 23.2 | 0.01 ± 0.01 | 0.05 ± 0.01 | 130 ± 40 | 0.003 ± 0.001 | 0.767 | S1 |
| A600t | t | AE | 6.4 | 0.9 ± 0.5 | 0.02 ± 0.01 | 3 ± 1 | 0.002 ± 0.001 | 0.948 | S1 |
| A600t | t | AE | 9.7 | 0.3 ± 0.4 | 0.05 ± 0.02 | 7 ± 3 | 0.003 ± 0.001 | 0.812 | S1 |
| A600t | t | AE | 13.3 | 0.6± 0.4 | 0.07 ± 0.01 | 5 ± 1 | 0.005 ± 0.002 | 0.921 | S1 |
| A600t | t | AE | 23 | 0.2 ± 0.1 | 0.13 ± 0.01 | 14 ± 3 | 0.005 ± 0.002 | 0.857 | S1 |
| exp | frac | var | $c_0$ [µM] | $k$ [1/h] | $I_p$ | $T_m$ [h] | $I_c$ | $R^2_{adj}$ | Fit eqn |
| ThTgain | t | HU | 5.3 | 0.5 ± 0.5 | 13 ± 4 | 3 ± 1 | 1.7 ± 0.6 | 0.761 | S1 |
| ThTgain | t | HU | 8.5 | 0.15 ± 0.1 | 20 ± 2 | 11 ± 3 | 1.4 ± 0.5 | 0.891 | S1 |
| ThTgain | t | HU | 12.6 | 0.06 ± 0.07 | 40 ± 10 | 30 ± 10 | 1.4 ± 0.5 | 0.918 | S1 |
| ThTgain | t | HU | 20.2 | 0.05 ± 0.02 | 50 ± 10 | 40 ± 10 | 1.2 ± 0.4 | 0.945 | S1 |
| ThTgain | t | HB | 6.9 | 0.02 ± 0.02 | 30 ± 10 | 50 ± 10 | 1.7 ± 0.6 | 0.575 | S1 |
| ThTgain | t | HB | 9 | 0.06 ± 0.06 | 40 ± 10 | 40 ± 10 | 1.2 ± 0.4 | 0.939 | S1 |
| ThTgain | t | HB | 13.9 | 0.03 ± 0.03 | 70 ± 20 | 80 ± 30 | 1.9 ± 0.4 | 0.938 | S1 |
| ThTgain | t | HB | 23.2 | 0.02 ± 0.03 | 60 ± 20 | 100 ± 40 | 0.9 ± 0.3 | 0.836 | S1 |
| ThTgain | t | AE | 6.4 | 0.9 ± 0.4 | 10 ± 3 | 3 ± 1 | 1.5 ± 0.5 | 0.896 | S1 |
| ThTgain | t | AE | 9.7 | 0.2 ± 0.2 | 18 ± 6 | 8 ± 3 | 2.1 ± 0.5 | 0.758 | S1 |
| ThTgain | t | AE | 13.3 | 0.02 ± 0.0 | 40 ± 10 | 60 ± 10 | 1.5 ± 0.5 | 0.754 | S1 |
| ThTgain | t | AE | 23 | 0.013 ± 0.003 | 140 ± 30 | 180 ± 90 | 2.3 ± 0.5 | 0.902 | S1 |
| ThTgain | t* | AE | 6.4 | 0.9 ± 0.4 | 10 ± 3 | 3 ± 1 | 1.5 ± 0.5 | 0.896 | S1 |
| ThTgain | t* | AE | 9.7 | 0.3 ± 0.5 | 17 ± 5 | 5 ± 2 | 1.5 ± 0.5 | 0.757 | S1 |
| ThTgain | t* | AE | 13.3 | 0.4 ± 0.4 | 20 ± 10 | 5 ± 2 | 1.5 ± 0.5 | 0.819 | S1 |

| | | | | | | | | | |
|---|---|---|---|---|---|---|---|---|---|
| ThTgain | t* | AE | 23 | 0.09 ± 0.05 | 40 ± 10 | 20 ± 6 | 1.4 ± 0.5 | 0.921 | S1 |
| exp | frac | var | $c_0$ [µM] | $k$ [1/h] | $I_p$ | $T_m$ [h] | $I_c$ | $R^2_{adj}$ | Fit eqn |
| A280s | s | HU | 8.5 | 0.04 ± 0.06 | 0.03 ± 0.01 | 30 ± 10 | 0.003 ± 0.001 | 0.741 | S2 |
| A280s | s | HU | 12.6 | 0.04 ± 0.04 | 0.06 ± 0.01 | 20 ± 10 | 0.006 ± 0.001 | 0.900 | S2 |
| A280s | s | HU | 20.2 | 0.02 ± 0.04 | 0.09 ± 0.03 | 50 ± 20 | 0.020 ± 0.004 | 0.681 | S2 |
| A280s | s | HB | 13.9 | 0.01 ± 0.03 | 0.05 ± 0.01 | 110 ± 50 | 0.010 ± 0.003 | 0.881 | S2 |
| A280s | s | HB | 23.2 | 0.01 ± 0.1 | 0.09 ± 0.03 | 210 ± 90 | 0.024 ± 0.008 | 0.672 | S2 |
| A280s | s | AE | 9.7 | 0.1 ± 0.2 | 0.039 ± 0.008 | 9 ± 5 | 0.001 ± 0.000 | 0.898 | S2 |
| A280s | s | AE | 13.3 | 0.1 ± 0.1 | 0.06 ± 0.01 | 14 ± 7 | 0.003 ± 0.001 | 0.841 | S2 |
| A280s | s | AE | 23 | 0.04 ± 0.05 | 0.10 ± 0.03 | 40 ± 20 | 0.009 ± 0.002 | 0.874 | S2 |
| exp | frac | var | $c_0$ [µM] | $k$ [1/h] | $I_p$ | $T_m$ [h] | $I_c$ | $R^2_{adj}$ | Fit eqn |
| RVgain | s | HU | 12.6 | 0.02 ± 0.02 | 1.4 ± 0.4 | 90 ± 50 | 0.4 ± 0.1 | 0.797 | S2 |
| RVgain | s | HU | 20.2 | 0.01 ± 0.01 | 2.2 ± 0.5 | 150± 70 | 0.6 ± 0.1 | 0.679 | S2 |
| RVgain | s | HB | 13.9 | 0.01 ± 0.01 | 2.2 ± 0.6 | 160± 50 | 0.8 ± 0.3 | 0.880 | S2 |
| RVgain | s | HB | 23.2 | 0.01 ± 0.04 | 4 ± 1 | 180± 90 | 0.4 ± 0.1 | 0.900 | S2 |
| RVgain | s | AE | 13.3 | 0.05 ± 0.04 | 2.2 ± 0.4 | 30 ± 10 | 0.4 ± 0.1 | 0.887 | S2 |
| RVgain | s | AE | 23 | 0.04 ± 0.05 | 4 ± 1 | 40 ± 20 | 0.7 ± 0.1 | 0.839 | S2 |

**Table S3.** Fit parameters of aggregation screen at various initial concentrations $c_0$ with parameter-adjusted R-squared $R^2_{adj}$ >0.5 ('adjrsquare' fit quality estimator of the MATLAB 'fit' function). Tabulated are experimental readout mode (exp), sample fraction (frac, 't' for total, 's' for supernatant after centrifugation),TTR variant (var), $c_0$ of the common stocks, $R^2_{adj}$, the used 'fit eqn', and the fitted parameters; fit parameters were the exponential fit rate $k$, the transition midpoint time $T_m$; the signal offset parameter $I_c$, and the initial intensity $I_0$, or the plateau intensity $I_p$ depending on which fit model was used. ($I_c$ and $I_p$ are unitless for normalized readouts ThT gain and RV gain, and given in arbitrary units of extinction for light extinction measurements). The ThT fluorescence intensity gain for aeTTR appears to be biphasic, and we included another fit in the table (t*) where the last datapoint (240h) was dropped for fitting to capture only the first plateau transition. The RV gain was allowed to go below RVgain=1, which was experimentally observed in some samples but is only possible due to dye fluorescence quenching artifacts.
